# A membrane-anchored inhibitor of papain-like cysteine proteases promotes *Pseudomonas* root colonization

**DOI:** 10.64898/2026.06.02.729523

**Authors:** Daniel Moser, Christian-Frederic Kaiser, Nina Solia, Lioba Rueger, Farnusch Kaschani, Brian C. Mooney, Ute Meyer, Athul Vijayan, Ulla Neumann, Renier A. L. van der Hoorn, Guido Grossmann, Tonni Grube Andersen, Gunther Doehlemann, Johana Misas Villamil

**Affiliations:** Institute for Plant Sciences, University of Cologne, Cologne, Germany; Institute for Cell and Interaction Biology, Heinrich-Heine University Duesseldorf, Duesseldorf, Germany; Max Planck Institute for Plant Breeding Research, Cologne, Germany; Analytics Core Facility Essen (ACE), University of Duisburg-Essen, Essen, Germany; The Plant Chemetics Laboratory, Department of Biology, University of Oxford, Oxford, UK; Cluster of Excellence on Plant Sciences (CEPLAS), Germany

**Keywords:** PLCPs, *Pseudomonas*, lipoprotein, surface-exposed, root colonization, commensal, chagasin inhibitor, OMVs

## Abstract

*Pseudomonas* species, spanning both beneficial and pathogenic lifestyles, possess conserved mechanisms to modulate plant immunity. Nevertheless, the mechanisms by which commensal bacteria establish and maintain host colonization remain poorly understood. Here, we report the characterization of a *Pseudomonas* chagasin-like protease inhibitor (Cpi1), conserved across pseudomonads representing a novel class of membrane-anchored PLCP inhibitor. Unlike previously described secreted protease inhibitors, *P. putida* Cpi1 is a lipoprotein localized to the bacterial surface and outer membrane vesicles (OMVs), positioning it to selectively inhibit immune-related papain-like cysteine proteases (PLCPs) during host interactions. Functional assays demonstrated inhibition of maize PLCP activity in the nanomolar range, while *cpi1* deletion and chagasin motif mutants exhibited significantly impaired early root colonization, particularly in the meristematic and elongation zones. Besides, lack of *cpi1* resulted in an altered structure of a maize root-associated synthetic community. We hypothesize that, Cpi1 may protect critical bacterial surface proteins from cleavage by inhibiting plant proteases and thereby modulate the release of MAMPs, dampening host immune responses. Moreover, the release of Cpi1 *via* OMVs could further extend its function within the root periphery and the apoplast. Together, our results uncover a conserved, membrane-anchored mechanism among pseudomonads for subverting plant immunity and establishing host-microbe interactions.

## Introduction

Plants interact with diverse microbial communities above- and belowground. The root microbiome includes pathogenic, mutualistic and commensal microbes, which are defined depending on the effect they have on plant fitness (Bai *et al*., 2022). Commensals can become pathogenic or mutualistic under certain conditions and the interaction with the plant can be regulated through root exudates or the plant’s immune system. The plant immune system recognizes microbes and regulates microbial colonization (Jones & Dangl, 2006). Pattern-triggered immunity (PTI) is activated by microbe-associated molecular patterns (MAMPs) detected by pattern-recognition receptors (PRRs) such as flagellin-derived flg22 and its receptor FLS2 (Chinchilla *et al*., 2006; Gómez-Gómez & Boller, 2000; Gómez-Gómez *et al*., 1999). PTI triggers signaling cascades, including calcium influx, reactive oxygen species (ROS) bursts, and defense compound production (Conrath *et al*., 2015).

Proteases regulate plant immunity by releasing immune peptides, degrading microbial proteins, or modulating defense components (Liu *et al*., 2024; Van Der Hoorn, 2008). Among them, papain-like cysteine proteases (PLCPs) act as hubs in plant immunity, being activated during infections (Misas-Villamil *et al*., 2016; Shindo & Van der Hoorn, 2008). Many pathogens modulate PLCP activity to suppress plant immunity. The oomycete *Phytophthora infestans* secretes the cystatin-like inhibitors EPIC1 and EPIC2B, which target tomato PLCPs such as PIP1, Rcr3, and C14, thereby interfering with host immune responses (Kaschani *et al*., 2010; Tian *et al*., 2007). Similarly, the fungal pathogen *Passalora fulva (Cladosporium fulvum)* produces Avr2, an effector that inhibits Rcr3 and PIP1 (Shabab *et al*., 2008). Additionally, the protist *Plasmodiophora brassicae* employs a cystatin-like effector (SSPbP53) to inhibit *Arabidopsis thaliana* XCP1, contributing to clubroot disease (Pérez-López *et al*., 2021). Pit2, a core effector of the pathogenic fungus *Ustilago maydis*, which acts as substrate-mimicking molecule, contains a conserved 14 amino acids motif (cMIP) that inhibits apoplastic maize PLCPs to suppress plant immunity (Doehlemann *et al*., 2011; Mueller *et al*., 2013; Misas Villamil *et al*., 2019). Notably, cMIP is conserved in plant-associated bacteria and is required to inhibit PLCPs (Misas Villamil *et al*., 2019). Another example is the bacterial pathogen *Pseudomonas syringae* pv. *tomato* DC3000, which uses C14-inhibiting protein 1 (Cip1), a chagasin-like inhibitor, to suppress the tomato immune protease C14 and PIP1 and promote virulence (Shindo *et al*., 2016). Chagasin is a potent PLCP inhibitor first identified in the protist parasite *Trypanosoma cruzi*, which regulates the activity of endogenous cruzipain, a PLCP essential for parasitic infection (Monteiro *et al*., 2001; Santos *et al*., 2005). Chagasin-like inhibitors, defined by their immunoglobulin fold and conserved inhibitory loops, have been found in other protozoa, bacteria, and archaea (Costa & Lima, 2016). While PLCP suppression is a common strategy among pathogens (Misas-Villamil *et al*., 2016; Shindo & Van der Hoorn, 2008), its role in beneficial interactions remains unclear. There are only few studies describing the inhibition of PLCPs in the interaction of beneficial microbes with plants. For example, the beneficial endophytic fungus *Epichloë festucae* suppresses PLCP activity during colonization (Passarge *et al*., 2021).

The interplay between plant immune responses and microbial recruitment is central to a healthy rhizosphere (Hacquard *et al*., 2017). While plants deploy PTI to regulate microbial colonization, they also create an inviting niche through root exudates (Bulgarelli *et al*., 2013). Particularly in the rhizosphere, root exudates play a key role in shaping microbial communities (Bulgarelli *et al*., 2013). These exudates, comprising sugars, amino acids, and secondary metabolites, play a major role in shaping microbial communities and attracting beneficial bacteria (Haichar *et al*., 2014; Rizaludin *et al*., 2021). A prevalence of bacterial genera such as *Pseudomonas* has been observed in disease-suppressive soils (Gómez Expósito *et al*., 2017). For example, *P. putida* AA7 was identified as keystone species within the maize root microbiome, where its presence strongly influenced the structure of a seven-member synthetic community (SynCom) (Niu *et al*., 2017). Pseudomonads also play key roles in plant growth promotion and disease suppression (Bulgarelli *et al*., 2013; Rivilla & Malone, 2023). In Arabidopsis *P. putida* enhances root biomass by promoting lateral root formation (Esparza-Reynoso *et al*., 2024) and in radish *P. putida* suppresses *Fusarium* wilt disease and induces systemic resistance (de Boer *et al*., 2003). Notably, pseudomonads can exhibit either pathogenic or beneficial traits, depending on their genomic adaptations (Xin *et al*., 2018). Beneficial strains use chemotaxis, biofilm formation, and antimicrobial competition to establish themselves in the rhizosphere (Espinosa-Urgel & Ramos-González, 2023; Zboralski & Filion, 2020). Some strains suppress PTI through the type III secretion system (T3SS)-deliver effectors, (Loper *et al*., 2012), whereas others avoid immune recognition by modifying surface molecules (Liu *et al*., 2018; Yu *et al*., 2019). Together, these findings highlight that modulation or evasion of plant immunity is a widespread strategy among rhizosphere-associated pseudomonads to facilitate root colonization and persistence within the microbiome.

Despite the emerging importance of microbial immune suppression during root colonization, the molecular mechanisms by which commensal rhizobacteria interfere with host PLCP activity and influence microbial community assembly remain poorly understood. Here, we investigated the role of *Pseudomona*s chagasin-like protease inhibitor 1 (Cpi1) in plant-microbe and microbe-microbe interactions during plant colonization. We demonstrate that Cpi1 is a lipoprotein widely conserved across pseudomonads, irrespective of their host or lifestyle. Microscopy and biochemical analyses show that Cpi1 localizes to the bacterial outer membrane and outer-membrane vesicles and is surface-exposed. *P. putida* AA7 Cpi1 inhibits plant PLCPs and plays a critical role in shaping the early structure of a synthetic bacterial community (SynCom). Finally, we show that the conserved chagasin motif of Cpi1 is required for early root colonization by *P. putida* AA7, consistent with a role for PLCP inhibition during bacterial establishment on plant roots.

## Results

### Chagasin-like protease inhibitors are predicted lipoproteins conserved in *Pseudomonas* species

To explore the conservation of the chagasin-like protease inhibitor 1 (Cpi1) in *Pseudomonas* species colonizing different hosts and with different life styles (pathogens or commensals), we conducted a BLAST search (Camacho *et al*., 2009) analysis of orthologs within the *Pseudomonas* genus. Cpi1 orthologs were found in all Pseudomonads regardless of host and lifestyle (Fig. 1A, Suppl. Data 1), including the human pathogen *Pseudomonas aeruginosa* PA01, the tomato pathogen *P. syringae* pv. *tomato* DC3000, the beneficial root colonizer *P. fluorescens* Pf0-1 or the commensal maize root colonizer *P. putida* AA7. These results confirm previous reports (Shindo *et al*., 2016) and suggest that Cpi1 orthologs contribute to conserved mechanisms of host-associated colonization across diverse Pseudomonas lifestyles.

**Figure 1.**
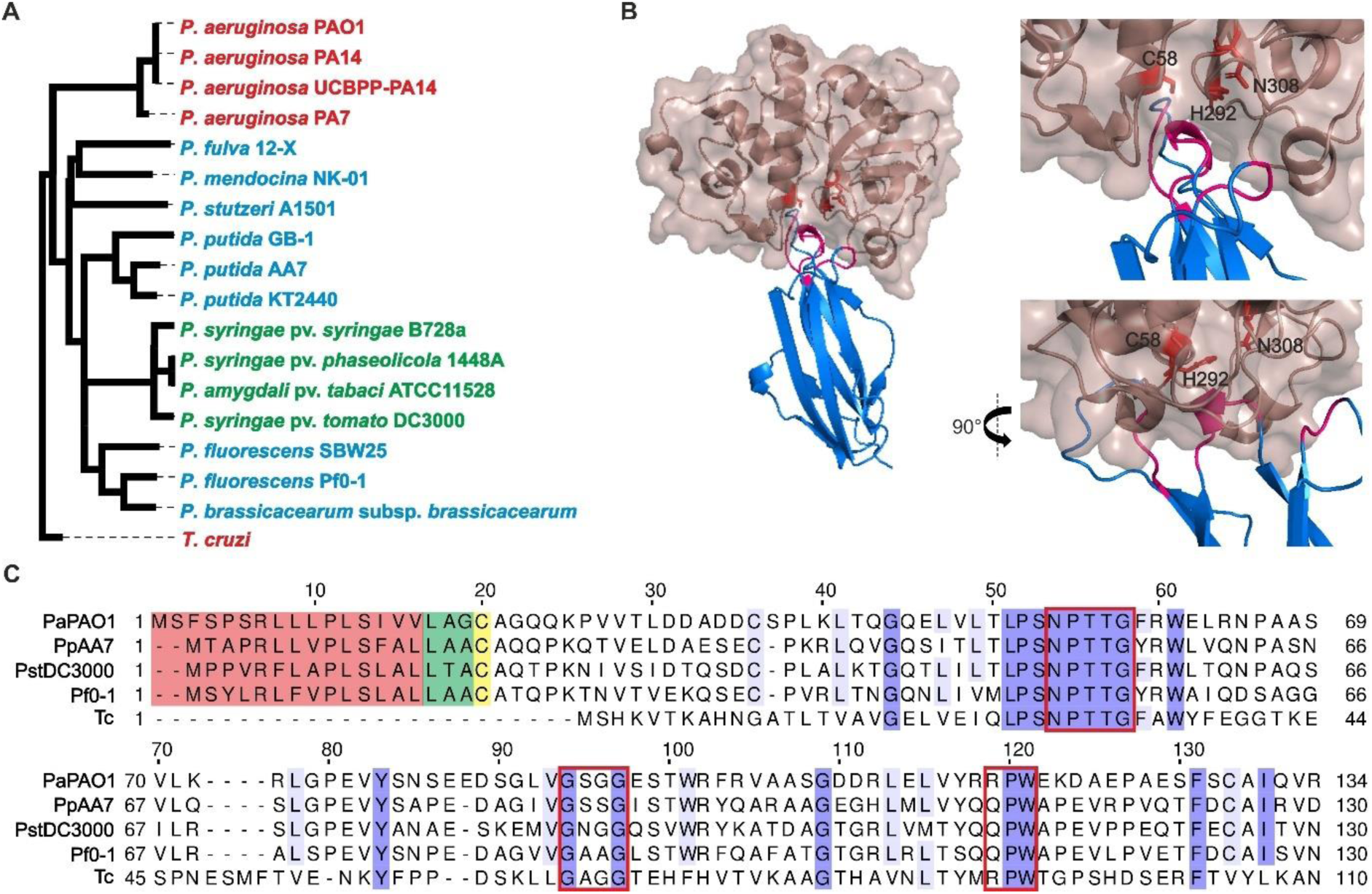
Chagasin-like inhibitors are ubiquitous lipoproteins in Pseudomonas. **(A) Phylogenetic tree of chagasin-like inhibitors of Pseudomonas species.** Chagasin-like inhibitors were identified in pseudomonads with different lifestyles (red: human pathogens, green: plant pathogens, blue: commensal bacteria) via DIAMOND BLASTp search. *Pseudomonas putida* AA7 (PpAA7) Cpi1 was used as a query for a search in the Pseudomonas database (Winsor *et al*., 2016). Identified orthologs (E < 0.001) and *Trypanosoma cruzi (T. cruzi)* chagasin were aligned and a phylogenetic tree was calculated with CLUSTAL Omega. **(B) Structural analysis of *P. putida* Cpi1 in complex with papain.** Left panel: Cpi1 (blue) is predicted to inhibit PLCPs (papain, dark red) by blocking the protease active site (red) with its chagasin motifs (pink). Cpi1 was modelled with AlphaFold3 (pTM = 0.77) without signal peptide. The model of Cpi1 was aligned to the structure of *T. cruzi* chagasin (root mean square deviation (RMSD): 1.04 Å) in a complex with papain (PDB: 3E1Z). Right panel: Zoom on loops with chagasin motif (pink) blocking active site of papain (red). Catalytic triad is labeled. **(C) Sequence alignment of chagasin-like inhibitors of different pseudomonads compared to *T. cruzi* chagasin.** Sequences of chagasin-like protease inhibitors (Cpi1) of *Pseudomonas aeruginosa* PAO1 (PaPAO1), *P. putida* AA7 (PpAA7), *Pseudomonas syringae* pv. *tomato* DC3000 (PstDC3000), *Pseudomonas fluorescence* Pf0-1 and *T. cruzi* (Tc) were aligned with CLUSTAL Omega. Red: signal peptide, green: lipobox, yellow: cysteine following the lipobox, light to dark violet: conserved amino acids with increasing conservation, red box: chagasin motifs.

AlphaFold3 predicted structures of several orthologs (Abramson *et al*., 2024) with the structure of *T. cruzi* chagasin (TcChagasin) in complex with the model PLCP papain (Redzynia *et al*., 2009) reveal a chagasin-like fold of Cpi1 orthologs of pseudomonads with either pathogenic, mutualistic or commensal lifestyles (Fig 1B, Fig S1). The Cpi1 ortholog of the maize rhizosphere keystone species *P. putida*, showed a highly similar strcture to chagasin (Fig. 1B, RMSD of 1.04 Å). In particular, the three loops containing the conserved chagasin motifs (NP[ST][ST]G, GxGG, and RP[WF]) required for PLCP inhibition are predicted to block the active site (Fig. 1B; catalytic triad consisting of C58, H292, N308), similar to the inhibition mechanism observed for chagasin (dos Reis *et al*., 2008; Shindo *et al*., 2016). Other *Pseudomonas* Cpi1 orthologs from *P. syringae* pv. *tomato* DC3000 (previously named Cip1, Shindo *et al*., 2016), *P. aeruginosa* PAO1 and *P. fluorescens* Pf0-1, showed high sequence similarity with conserved chagasin motifs similar to *T. cruzi* (Fig. 1C) and have a conserved structural fold to block the active site of papain (Suppl. Fig. 1A-B). These conserved structural features of chagasin orthologs in *Pseudomonas* indicate a conserved function in inhibiting PLCPs.

Interestingly, *Pseudomonas* Cpi1-like proteins have a conserved signal peptide which is not present in *T. cruzi* chagasin (Fig. 1C). Signal peptide analysis of all identified orthologs using SignalP6 (Teufel *et al*., 2022) revealed that all except for two *Pseudomonas* chagasin-like proteins are predicted to be lipoproteins (Suppl. Fig. 1). Analyzed Cpi1 orthologs have a signal peptide for the secretory (Sec) pathway and a three amino acids lipobox that predicts processing as lipoproteins (LoVullo *et al*., 2015; Natale *et al*., 2008; Teufel *et al*., 2022) (Fig. 1C).

During periplasmic processing in Gram-negative bacteria, the lipobox containing signal peptide is removed and a diacylglyceryl moiety as well as another acyl group are attached to the new N-terminal cysteine (C18), anchoring the mature lipoproteins to the inner membrane (Gupta *et al*., 1993; Sankaran & Wu, 1994; Tokunaga *et al*., 1982, 1984). From the inner membrane, lipoproteins can also be peripherally attached to the inner or outer leaflet of the outer membrane facing the periplasm or the extracellular space respectively, however, the exact processes involved in their translocation are not yet well characterized (Cole *et al*., 2021). The presence of the lipobox in Cpi1 suggests that this chagasin-like inhibitor is membrane-anchored.

To investigate the localization and biological function of *Pseudomonas* Cpi1 proteins, we employed the commensal strain *P. putida* AA7 (hereafter *P. putida*). Marker-free gene deletion mutants (Δ*cpi1*) were generated (Suppl. Fig. 2A-F). Two independent *P. putida* mutants (Δ*cpi1_*#2 and #3) did not show a growth defect when grown in full medium (Suppl. Fig. 2F). Δc*pi1* was genetically complemented using an episomal vector with the c*pi1* gene and the addition of a C-terminal 3xFlag-tag resulting in Δ*cpi1-cpi1-flag*. The *P. putida* strains WT, Δ*cpi1* and Δ*cpi1*-*cpi1-flag* grown in full medium were tested for *cpi1* expression after one and five hours. The absence of *cpi1* transcripts was confirmed in the Δ*cpi1* mutant, while the WT strain expressed *cpi1* in axenic cultures. Notably, expression of *cpi1* was similar when bacteria were incubated in M9 minimal medium (Suppl. Fig. 2J), indicating that *Pseudomonas cpi1* is expressed in axenic cultures independent of the nutrient status of the medium. The Δ*cpi1*-*cpi1-flag* complementation strain showed a strong overexpression of *cpi1* compared to the WT (approx. 100-fold; Suppl. Fig. 2J).

### Cpi1 localizes to the bacterial outer membrane

To investigate whether *Pseudomonas* chagasin-like proteins are indeed membrane-anchored lipoproteins, fluorescence microscopy was performed. The gene encoding cytosolic superfolder GFP (sfGFP) was stably integrated into the *P. putida* genome to generate WT-sfG (Suppl. Fig. 2G–I). An episomal vector enabling constitutive expression of mCherry alone or Cpi1-mCherry was subsequently introduced into the WT-sfG strain, resulting in WT-sfGmC and WT-sfG-Cpi1-mC. Fluorescence microscopy showed that Cpi1-mCherry was significantly enriched at the cell periphery and its signal is reduced in the center of the cells compared to cytosolic sfGFP and mCherry signals (Fig. 2A-B), indicating that Cpi1-mCherry is localized at the bacterial membrane.

**Figure 2.**
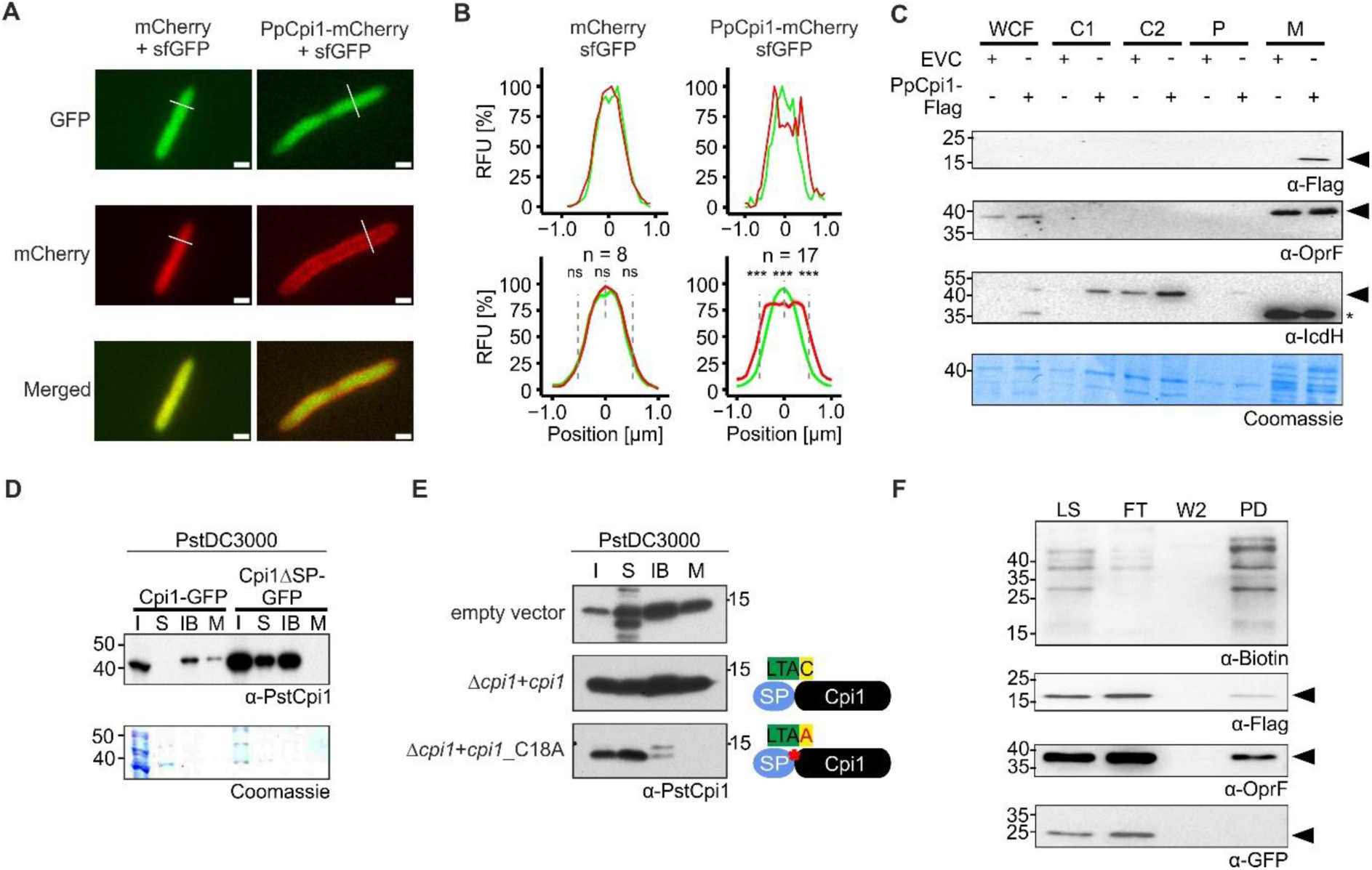
Cpi1 is localized at the surface of the bacterial outer membrane. **(A) Cpi1-mCherry fusion protein localizes to the cell envelope of *P. putida*.** Representative microscopy images of *P. putida* WT-sfG expressing mCherry (left) or Cpi1-mCherry (right). **(B) Quantification of fluorescence signals.** Fluorescence intensity histograms of WT-sfG cells expressing mCherry (left) or Cpi1-mCherry (right) shown in (A). Lower panel: Mean relative fluorescence intensity profiles of different WT-sfG cells (n). The mean is shown as a dark line with SEM indicated by shading. Differences between sfGFP and mCherry signals at −0.52, 0 and 0.52 μm from the cell centre were assessed using paired t-tests. Significance is indicated as ***P<0.001, **P<0.01, *P<0.05, ns (not significant). **(C) Cpi1 is detected in the membrane fraction**. Cells of *P. putida* Cpi1-Flag were fractionated (WCF: whole cell fraction, C1-C2: Cytosol, P: Periplasm, M: Membrane). Western blot analysis of Cpi1-Flag (approx. 17 kDa), a known outer membrane protein porin F (OprF) (approx. 36 kDa) and the cytosolic marker isocitrate dehydrogenase (IcdH) (approx. 45 kDa) was performed. Asterisk indicates background signal. **(D) Membrane localization of *P. syringae* Cpi1 is dependent on its signal peptide.** Cells of *P. syringae* DC3000 (PstDC3000) overexpressing Cpi1-GFP and a deletion mutant without signal peptide Cpi1ΔSP-GFP were fractionated (I: Input sample, S: Soluble fraction, IB: Inclusion bodies, M: Membrane fraction). Western blot analysis was performed to monit r the localization of Cpi1-GFP (approx. 40 kDa) using PstCpi1-specific antibodies. **(E) C18 in lipobox is required for membrane localization.** *P. syringae* DC3000, the complementation strain Δ*cpi1*+Cpi1 and the mutant strain Δ*cpi1*+Cpi1_C18A were fractionated (I: Input sample, S: Soluble fraction, IB: Inclusion bodies, M: Membrane fraction). Western blot analysis was performed to monitor Cpi1 (approx. 14 kDa) using PstCpi1-specific antibodies. **(F) Cpi1 is localized at the surface of the bacterial outer membrane**. Biotin labeling using the membrane-impermeable biotinylation reagent Sulfo-NHS-Biotin was used to covalently label all surface-localized proteins of the WT-sfG strain expressing Cpi1-Flag (LS: labeled surface proteins). After labeling, the Sulfo-HNS-Biotin biotinylated proteins were pulled down using streptavidin beads (FT: flow-through, W2: 2^nd^ wash fraction, PD: pull-down). Proteins were detected via Western Blot using specific antibodies: α-Biotin to monitor biotinylated proteins, α-Flag for Cpi1-Flag, α-OprF for the outer membrane protein F (OprF) as a positive control and α-GFP for the cytosolic sfGFP as a negative control. All these experiments were performed with at least three independent biological replicates.

To confirm the fluorescence microscopy analysis, we followed a biochemical approach using a Cpi1-Flag strain. We separated cell compartments (Paredes-Osses & Hardie, 2013) and used antibodies against proteins with known localization in other bacterial strains as controls to confirm the successful cell fractionation. We observed that Cpi1-Flag (approx. 17 kDa) was found only in the membrane fraction, similar to the known membrane marker protein “outer membrane protein porin F” (OprF) (approx. 36 kDa) (Fig. 2C). In contrast, the cytosolic marker protein “isocitrate dehydrogenase H” (IcdH) (approx. 45 kDa) was found only in the cytosolic fractions. The biochemical analysis together with the fluorescence microscopy indicate that Cpi1 is a membrane-anchored lipoprotein in *P. putida*.

To test whether other Cpi1 orthologs are also lipoproteins, cell fractionation experiments with the plant pathogen *P. syringae* pv. *tomato* DC3000 (PstDC3000) were performed. Western blot analysis of the cell fractions using a specific PstCpi1 antibody (Shindo *et al*., 2016) of the strain overexpressing Cpi1-GFP showed the presence of Cpi1 in the insoluble and membrane fraction. In contrast, the mutant strain lacking the signal peptide (Cpi1ΔSP-GFP) showed a signal in the soluble fraction, indicating Cpi1 secretion and that the signal peptide functions as a membrane-anchor signal (Fig. 2D). We further tested if the conserved cysteine C18 in the lipobox is required for the observed membrane localization of Cpi1 by exchanging C18 to A18 in the strain Δ*cpi1*-Cpi1_C18A. Strikingly, *P. syringae* Cpi1 was absent in the membrane fraction in PsCpi1_C18A, but not in the WT or complementation strain

Δ*cpi1*_*cpi1*, indicating that the conserved cysteine is necessary and required for the observed membrane localization of Cpi1 (Fig. 2E). In conclusion, these results show that both in *P. putida* and *P. syringae* Cpi1 is a membrane-anchored lipoprotein, and the conserved cysteine in the lipobox is required for membrane localization.

### Cpi1 is a surface-exposed inhibitor of plant PLCPs

Chagasin-like inhibitors have been reported to inhibit only C1 family proteases (Costa & Lima, 2016). Similarly, *P. syringae* Cpi1 (former Cip1) was shown to inhibit tomato PLCPs (Shindo *et al*., 2016). Since Cpi1 was found to be membrane-anchored, we tested whether Cpi1 is surface-localized, which would be a prerequisite to interfere with host-derived PLCPs. Therefore, biotin labeling of surface-exposed proteins of *P. putida* WT-sfG-Cpi1-Flag (overexpressing *cpi1* and genomic *sfgfp*) was performed.

Cell surface proteins were covalently labeled using the membrane-impermeable Sulfo-NHS-Biotin reagent (Daniels & Amara, 1998; Huh & Wenthold, 1999) and biotinylated proteins from lysed cells were pulled down using streptavidin beads. Streptavidin-HRP blots showed proteins of different sizes indicating a successful enrichment of biotinylated proteins (Fig. 2F). To validate specific labeling of surface proteins, antibodies against OprF and sfGFP were employed. The outer membrane protein OprF was successfully labeled and detected, while the cytosolic sfGFP was not detected (Fig. 2F), indicating that the biotinylation method is reliable for surface protein labeling and that bacterial cells remained intact during the procedure. Additionally, Flag antibodies revealed Cpi1 enrichment in the biotinylated fraction, further confirming Cpi1-Flag is both, membrane-bound and surface-localized.

Gram-negative bacteria are known to produce outer membrane vesicles (OMVs) that encapsulate a variety of cargos, such as nucleic acids and proteins, which contribute to intercellular signaling between bacteria, the surrounding environment, and host organisms (Rudnicka *et al*., 2022). Thus, the membrane localization of Cpi1 suggests a potential association with OMVs. To determine whether *P. putida* Cpi1 localizes to OMVs, vesicles were purified from WT and Cpi1-Flag strains and their quality and abundance were evaluated using transmission electron microscopy (Fig. 3A). The diameter of purified vesicles from both strains ranged from 55 to 80 nm, which is consistent with the size reported for *P. putida* OMVs (Bitzenhofer *et al*., 2024; Salvachúa *et al*., 2020). Purified vesicles were further analyzed by Western blot. Cpi1-Flag was detected exclusively in the Cpi1-Flag sample, whereas OprF, used as a positive control, was detected in both WT and Cpi1-Flag preparations confirming the presence of Cpi1 in *P. putida* OMVs (Fig. 3B).

**Figure 3.**
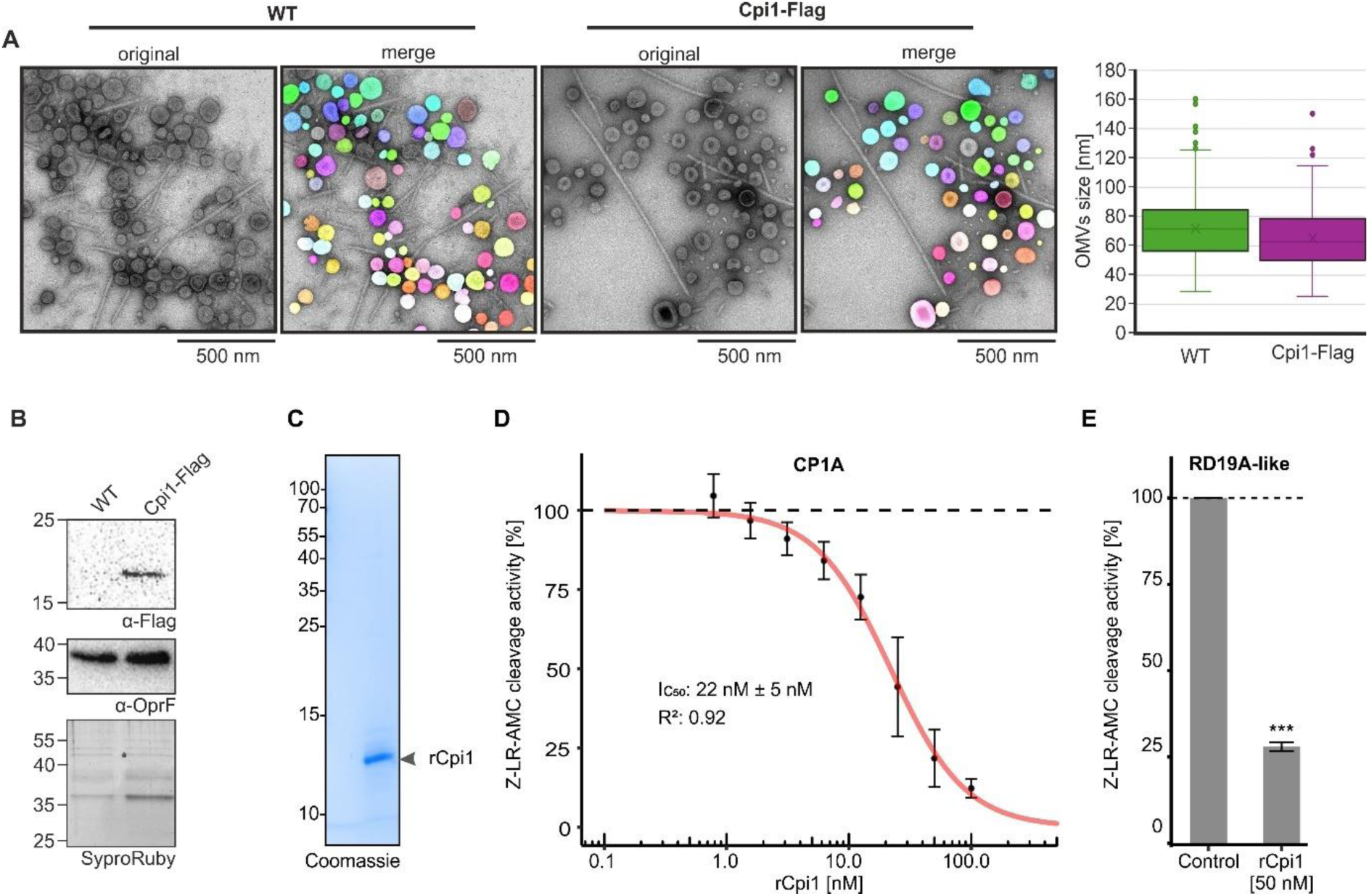
Cpi1 is present in OMVs and inhibit maize PLCPs. **(A**) **Imaging and quantification of outer membrane vesicles (OMVs)**. OMVs from WT and Cpi1-Flag overexpression strains were isolated by density gradient centrifugation and analyzed by electron microscopy. Representative micrographs and corresponding segmented images are shown; segmentation was curated to remove false positives and edge artifacts. Scale bar, 500 nm. OMV diameter was quantified from ten micrographs per genotype (WT, n= 936; Cpi1-Flag, n = 521) and is presented as box-and-whisker plots (boxes: IQR; median: line; mean: cross). **(B) Cpi1 is detected in OMVs.** Western blot analysis of isolated OMVs were performed with the α-Flag antibodies to detect Cpi1-Flag and the OprF antibody, a common membrane protein as a control. SYPRO Ruby staining is shown as a loading control. **(C) Production of recombinant *P. putida* Cpi1**. His-SUMO-Cpi1 without signal peptide was heterologous expressed in *E. coli* and purified by affinity chromatography. Afterward, the His-SUMO tag was removed and tag-free rCpi1 was further purified by size-exclusion chromatography. Purity of rCpi1 (12.6 kDa) was confirmed by SDS-PAGE followed by Coomassie staining. **(B) Cpi1 inhibits the activity of the apoplastic immune protease CP1A**. Heterologously expressed CP1A in *N. benthamiana* was pre-incubated with increasing concentrations of rCpi1. The activity of CP1A was measured using the substrate Z-LR-AMC. The Z-LR-AMC cleavage activity of each treated sample was normalized to the water control set to 100%. IC_50_ was estimated by fitting the data with the sigmoid Hill function. This experiment was performed with three independent biological replicates with each having at least two technical replicates. Means are represented as black dots. Error bars represent SEM. **(C) Cpi1 inhibits the activity of the root immune protease RD19A-like**. RD19A-like heterologously expressed in *N. benthamiana* was preincubated with either water or 50 nM rCpi1. The activity of RD19A-like was measured with the substrate Z-LR-AMC where the increase of AMC fluorescence was monitored over time as described in (B). Means are represented as grey bars and SEM are shown. A two-sided t-test was performed (P<0.05: *; P<0.01: **; P<0.001: ***). All experiments were repeated at least three times with similar results.

To test whether Cpi1 is an inhibitor of root PLCPs, Cpi1 lacking its signal peptide (SP) was recombinantly expressed in *E. coli* (Fig. 3C). The recombinant protein (rCpi1) was tested for inhibitory activity of maize PLCPs in a substrate cleavage assay. To this end, two immune-related maize PLCPs, ZmCP1A (van der Linde *et al*., 2012) and ZmRD19A-like, both active in the maize root apoplast (Schulze Hüynck *et al*., 2019), were heterologously expressed in *N. benthamiana* using *Agrobacterium-*mediated transformation. Apoplastic fluids containing recombinant ZmCP1A were then incubated with different concentrations of rCpi1. ZmCP1A was inhibited in a concentration-dependent manner resulting in an IC_50_ of 22 nM ± 5 nM (Fig. 3D, Suppl. Fig. 3). This IC_50_ value correlates with the pico- to nanomolar K_i_ reported for chagasin as well as *P. syringae* and *P. aeruginosa* chagasin-like inhibitors (dos Reis *et al*., 2008; Monteiro *et al*., 2001). Similarly, the activity of ZmRD19A-like was reduced to 75%, when inhibited by 50 nM rCpi1 (Fig. 3E, Suppl. Fig. 3). These data demonstrate that Cpi1 is an inhibitor of maize immune-related PLCPs.

### Cpi1 promotes early root colonization in maize and *Arabidopsis*

To investigate the role of Cpi1 during bacterial community interactions, we performed SynCom colonization assays of maize plants grown for up to 14 days in sterile soil using a 7-member maize root bacterial community (Niu *et al*., 2017). We compared communities containing either *P. putida* WT, or the Cpi1 deletion mutant (Δ*cpi1*). At different time points (3 hpi, 7 dpi and 14 dpi) the rhizoplane was harvested and the community structure was determined by 16S DNA amplicon sequencing (Fig. 4A). We observed that the community structure in the SynCom with Δ*cpi1* significantly changed after 3 hpi compared to the SynCom with WT, whereas at later time points the observed early differences were less marked, reflecting and increasing stabilization of the community over time (Fig. 4B, Suppl. Fig. 4). The relative abundance of *E. cloacae*, another keystone species in the community besides *P. putida*, increased in the absence of Cpi1, whereas the abundance of *C. indologenes*, *H. frisingense* and *P. putida* itself decreased after 3hpi (Fig. 4C, Suppl. Fig. 4). At 7 dpi *C. indologenes* was significantly more abundant in the community in the absence of Cpi1, whereas *H. frisingense* was less abundant (Fig. 4C, Suppl. Fig. 4). At 14 dpi *C. pusillum* was more abundant in the absence of Cpi1 while the abundance of *E. cloacae* was reduced (Fig. 4C, Suppl. Fig. 4). Notably, *P. putida* WT was more abundant than *P. putida* Δ*cpi1* only after 3 hpi but not at later time points, which might indicate an effect of Cpi1 in early root colonization (Fig. 4D). This is consistent with previous reports indicating that the initial phase of *P. putida* colonization is highly active, with the bacterial population reaching its maximum relative to root biomass 24–48 h after seedling inoculation (Espinosa-Urgel *et al*., 2002). In conclusion, significant Cpi1-dependent changes in the root community structure were observed at 3 hpi and 7 dpi demonstrating that Cpi1 has an effect on the SynCom structure, particularly at early stages of root colonization. One possible explanation is that Cpi1 directly affects community structure by inhibiting bacterial C1 family cysteine proteases (PLCPs). To test whether SynCom members encode these enzymes, we performed a BLASTp search using C1 family protease sequences from the MEROPS database as queries (Altschul *et al*., 1990; Camacho *et al*., 2009; Rawlings *et al*., 2014). Only one putative C1 family peptidase (Cin_2970) (E<10^-3^) was identified in *C. indologenes*, while no PLCPs were found in the other bacterial genomes. Therefore, we consider it more likely that the observed early Cpi1-dependent shifts in SynCom structure result from inhibition of plant PLCPs rather than interbacterial PLCP inhibition.

**Figure 4.**
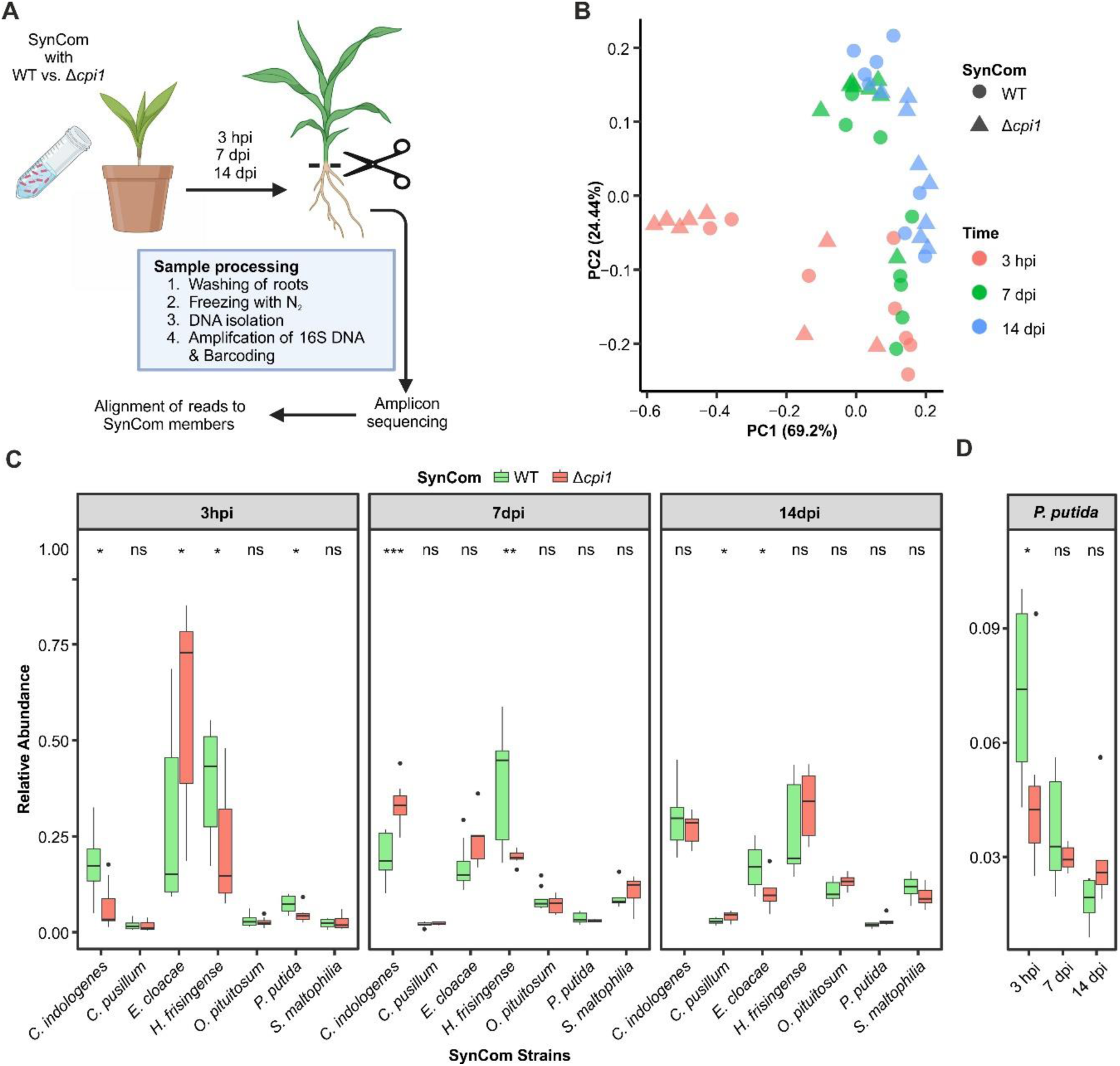
Cpi1 modulates the structure of a maize synthetic community (SynCom). **(A) Schematic representation of the experimental design.** Maize plants were grown in soil containing the six SynCom members and with either *P. putida* WT or *P. putida*Δ*pci1* (SynCom inoculated to an OD_600_ of 0.0001). Maize roots were harvested at 3 hpi, 7 dpi and 14 dpi, washed several times by vortexing in water, and samples were frozen with liquid nitrogen. DNA was isolated and 16S sequence was amplified via PCR, barcoded and sent for amplicon sequencing. Reads were aligned to the SynCom members 16S RNA sequences and quantified. This experiment was performed once with eight biological replicates per timepoint and SynCom. Created in BioRender. Doehlemann, G. (2025) https://BioRender.com/knymui1. **(B) Principal Coordinate Analysis (PCoA) of quantified reads**. PCoA was performed using a Bray-Curtis dissimilarity matrix to graphically represent all samples. The principal components (PC1 and PC2) represent the variance in the data. Goodness of fit: PC1: 0.80%, PC2: 0.86%. **(C) Abundance of SynCom members over time**. The abundance of all SynCom members with *P. putida* WT (green) or Δ*pci1* (red) was quantified based on the aligned reads and normalized to the total amount of reads in a sample. A two-sided t-test was performed (P<0.05: *; P<0.01: **; P<0.001: ***). (D) PpPci1 promotes early root colonization of *P. putida*. The abundance of *P. putida* WT (green) or Δ*pci1* (red) was quantified based on the aligned reads and normalized to the total amount of reads in a sample. A two-sided t-test was performed (P<0.05: *; P<0.01: **; P<0.001: ***).

To investigate the role of Cpi1 in early root colonization a modified RootChip-8S (Denninger *et al*., 2019; Guichard *et al*., 2020) was used. *Arabidopsis thaliana* Col-0 roots were grown for 10 days into linear channels of a sterile chip (Kaiser *et al*., 2026). Roots were subjected to fluorescence imaging one day after inoculation with sfGFP-expressing *P. putida* strains: WT, Δ*cpi1*, the complemented strain Δ*cpi1-cpi1*, and Δ*cpi1-NPTTG^mut^*, lacking the inhibitory chagasin domain. Bacterial coverage was calculated as the percentage of root-associated area occupied by GFP fluorescence after subtraction of root autofluorescence. GFP signals were mostly observed within the first 1500 µm from the root tip (Fig. 5A-B, Suppl. Fig. 5), indicating that the bacteria mostly colonize the younger parts of the root. Within this area, GFP fluorescence was significantly reduced for Δc*pi1* compared to WT (Fig. 5A-B, Suppl. Fig. 5), indicating that Cpi1 is required for efficient root colonization (Fig. 5A-B). To further dissect spatial differences in root colonization, developmental zones were assigned based on Verbelen *et al*. (2006) (Fig. 5A), and GFP fluorescence per µm was quantified for each zone. Compared to WT, Δ*cpi1* showed significantly reduced GFP fluorescence in the meristematic, transition, and elongation zones, whereas fluorescence levels in the differentiation zone were comparable to WT (Fig. 5A–C). These results suggest that Cpi1 contributes to efficient colonization of the root tip and adjacent developmental zones. The complemented Δ*cpi1*-c*pi1* strain showed an increased colonization phenotype compared to Δ*cpi1*, confirming that Cpi1 is required for early root colonization (Fig. 5A-C). To test whether the early colonization phenotype is due to the inhibitory capacity of Cpi1 we generated a fluorescent chagasin mutant strain by deleting the chagasin motif (NPTTG) required for Cpi1 function (Shindo *et al*., 2016). Similar to the Δ*cpi1* strain, the chagasin mutant Δ*cpi1*-NPTTG*^mut^* showed significantly reduced colonization of the meristematic, transition and elongation zone in comparison to the WT (Fig. 5A-C). This result indicates that the inhibitory function of Cpi1 is necessary and required for *P. putida* early root colonization. In conclusion, Cpi1 modulates the early colonization of both *A. thaliana* and maize roots. The differential spatiotemporal *P. putida* colonization on *A. thaliana* indicates that Cpi1 supports early root colonization in the meristematic, transition and elongation zones by inhibiting plant PLCPs.

**Figure 5:**
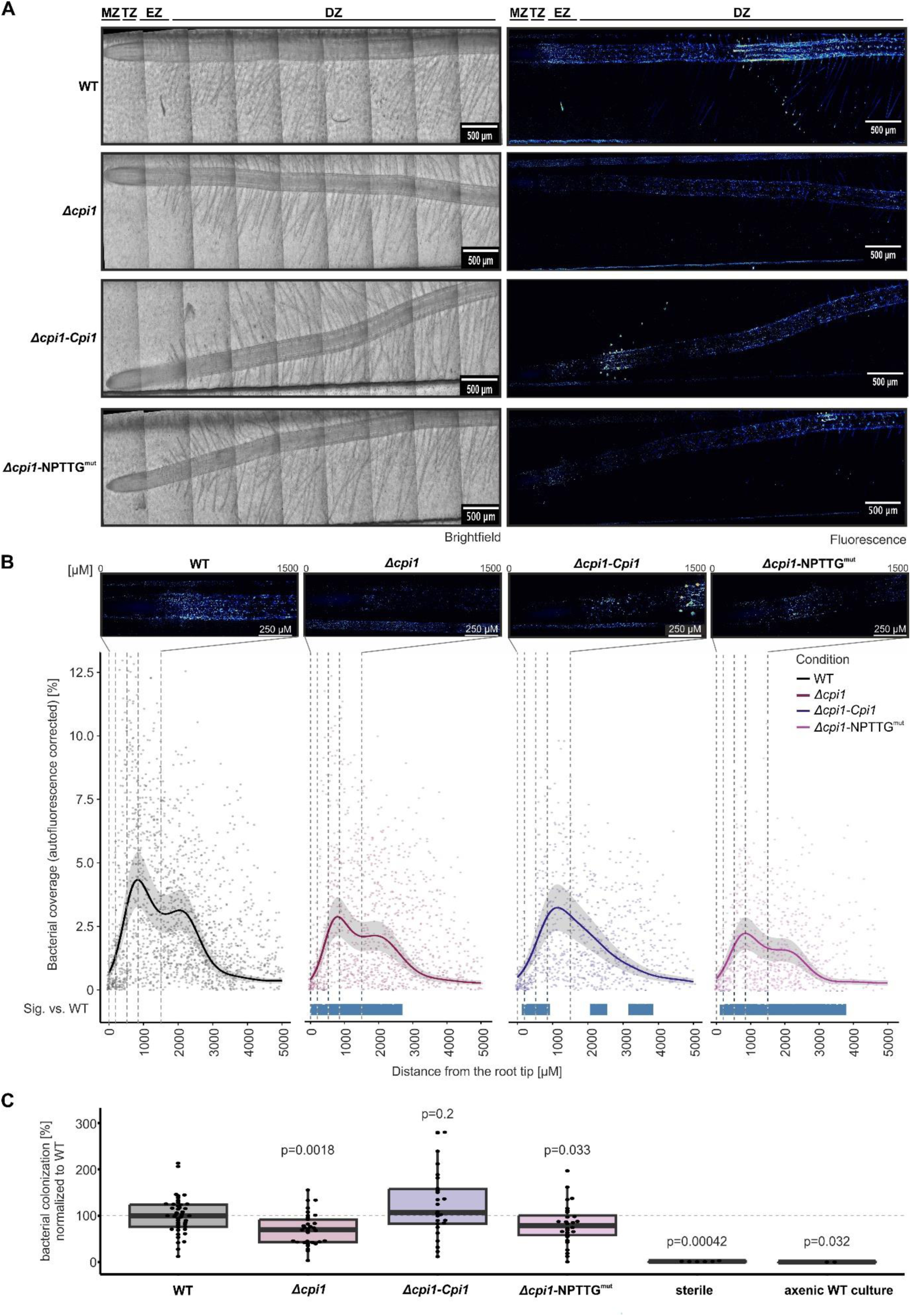
Cpi1 inhibitory function is required for early root colonization. **(A) Fluorescence microscopy of *A. thaliana* Col-0 roots colonized by *P. putida*.** *P. putida* strains expressing sfGFP: WT, Δ*cpi1*, Δ*cpi1-cpi1* and Δ*cpi1-NPTTG^mut^* were inoculated onto 10-day-old *A. thaliana* Col-0 roots grown in a RootChip8NF. GFP fluorescence was imaged 1 dpi. Fluorescence intensity is shown using the royal lookup table with identical intensity limits across images (low: blue, high: red). Each panel shows a representative root from the same imaging batch; images were stitched from 18 fields of view and background corrected. Root zones are indicated on the top: meristematic zone (MZ, 0–200 µm from the root tip), transition zone (TZ, 200–520 µm), elongation zone (EZ, 520–850 µm), differentiation zone (DZ, >850), estimated according to Verbelen *et al*. (2006). Scale bar: 500 µm. **(B) Cpi1 promotes root colonization of the meristematic, transition and elongation zones**. Fluorescence intensity was used to segment bacterial clusters (≥ 3.5 µm²). The segmented area was summed within 200 × 100 µm windows centered on the root midline and normalized to obtain bacterial surface coverage. Colonization profiles along the root were modeled using generalized additive models (GAMs), with coverage as the response and distance from the root tip as a smooth term. A position-dependent baseline from non-inoculated controls was subtracted prior to modeling (S5A-B). Condition-specific smooths were fitted with imaging date and image identity as random effects. Solid lines indicate fitted GAMs, shaded regions show standard error, and points represent individual measurements. Regions significantly different from WT are indicated below the plot (blue). The image above shows the cropped region (0–1500 µm from the root tip). **(C) Colonization of the root apical meristem requires Cpi1 inhibitory function**. Fluorescence signal was used to segment bacterial clusters as described in (B). Segmented bacterial area was summed within a 200 µm-wide window centered on the root midline and normalized to the corresponding root area (Distance to the root tip < 1500 µm) to obtain bacterial surface coverage per root. Each root was normalized to the median wild-type coverage within the same replicate (set to 0) to calculate the relative difference from WT colonization. Data are shown as box plots with individual roots displayed as jittered points. Significant differences from WT reference were assessed using two-sided Wilcoxon rank-sum tests, with p-values indicated above the box plots. Sample sizes: WT (N = 46 roots, 8 days), Δ*cpi1* (N = 33 roots, 6 days), Δ*cpi1*-c*pi1* (N = 26 roots, 5 days), Δ*cpi1-NPTTG^mut^* (N = 28 roots, 6 days), axenic WT culture and sterile roots (N = 3 roots, 1 day).

In summary, we show that Cpi1 inhibited plant proteases and promoted early root colonization in maize and Arabidopsis. In *P. putida*, Cpi1 contributed to protease inhibition, enhanced root colonization, and affected microbial community composition.

## Discussion

Structural analyses showed that Cpi1 and related pseudomonad proteins retain the key inhibitory motifs of canonical chagasins, including the loops occluding target protease active sites (dos Reis *et al*., 2008; Redzynia *et al*., 2008, 2009). Consistent with this conservation, Cpi1 inhibited maize PLCPs at nanomolar concentrations similar to other chagasin family members (dos Reis *et al*., 2008; Fu, 2009; Monteiro *et al*., 2001; Shindo *et al*., 2016), supporting a common evolutionary origin of bacterial and protozoal chagasin-like proteins (Sanderson *et al*., 2003).

Our study identified the chagasin-like inhibitor Cpi1 as a critical factor for early root colonization by *P. putida*. Unlike previously characterized PLCP inhibitors secreted to the extracellular space, such as EPIC1, Avr2 or Pit2 (Doehlemann *et al*., 2011; Kaschani *et al*., 2010; Misas Villamil *et al*., 2019; Mueller *et al*., 2013; Shabab *et al*., 2008; Tian *et al*., 2007) Cpi1 is a membrane-anchored lipoprotein that localizes to the bacterial cell surface and OMVs. In contrast to the regulation of the endogenous cruzipain by *T. cruzi* chagasin (Monteiro *et al*., 2001; Santos *et al*., 2005), we hypothesize that *Pseudomonas* chagasin-like inhibitors modulate the activity of plant PLCPs, as *P. putida* has no endogenous C1 family proteases and Cpi1 strongly inhibits apoplastic maize PLCPs. The membrane-anchored nature of Cpi1 suggests a spatially restricted mode-of-action distinct from secreted effectors. Reports of bacterial lipoprotein protease inhibitors are rare. The *Brucella abortus* lipoprotein Omp19 and the periplasmic inhibitor Ecotin protect bacterial surface proteins against host-derived proteases (Chung *et al*., 1983; De Meyer & Carlier, 2023; Eggers *et al*., 2004; McGrath *et al*., 1991, 1994; Pasquevich *et al*., 2019; Yang *et al*., 1998), illustrating how bacteria modulate proteolytic activity at the cell envelope. However, membrane-anchored chagasin-like inhibitors remain poorly characterized, and Cpi1 represents the first example implicated in PLCP inhibition.

To understand how Cpi1 promotes bacterial host colonization, it will be instrumental to explore the mechanism by which they interfere with the host. Due to its surface-exposed nature, one could hypothesize that Cpi1 protects surface-localized proteins or the cargos of the OMVs from PLCP-mediated cleavage (Fig. 6), similar to the protective functions described for Omp19 and Ecotin (Pasquevich *et al*., 2019; Eggers *et al*., 2004). Indeed, PLCPs have been implicated in the cleavage of an outer membrane protein (OMP) of the citrus bacterial pathogen *Candidatus liberibacter* causing the citrus greening disease Huanglongbing. OMP cleavage has been hypothesized to impair bacterial growth and/or to activate plant immune responses (McClelland *et al*., 2026). Such localized inhibition by Cpi1 could preserve proteins involved in motility, adhesion, or nutrient acquisition while limiting immune activation during early root colonization (Wadhwa & Berg, 2022). This spatially restricted inhibition could allow the bacterium to fine tune the suppression of plant immunity without broadly compromising the plant’s ability to respond to microbes. Our results support the notion that by inhibiting PLCPs, Cpi1 might also dampen early immune responses, e.g. by reducing the release of MAMPs, thereby facilitating bacterial colonization (Fig. 6).

**Figure 6:**
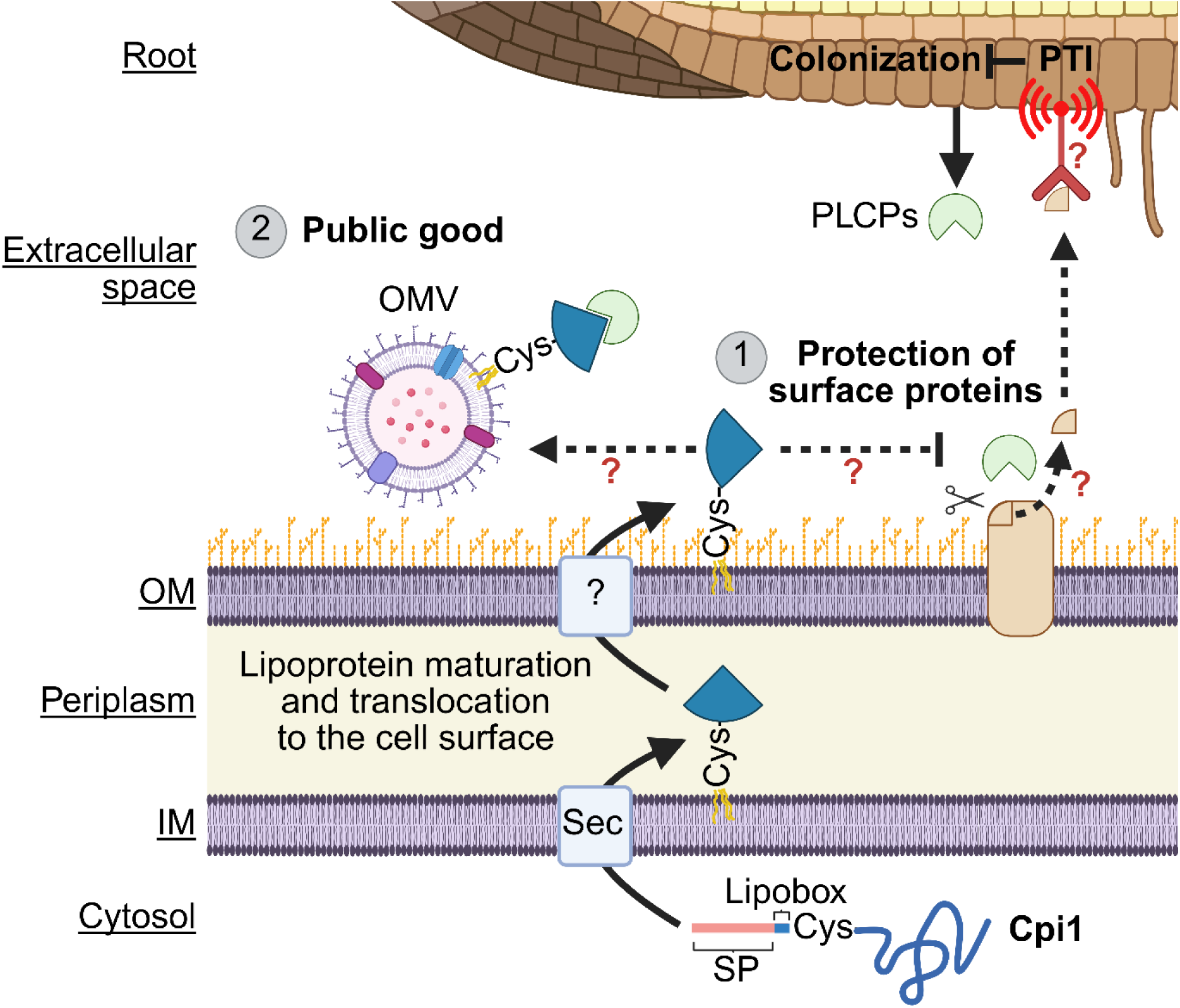
Model of Cpi1 modulation of root colonization in *P. putida*. The root commensal *P. putida* produces the chagasin-like protease inhibitor Cpi1. Cpi1 showed a novel mode of action for inhibitor delivery: Cpi1 is a predicted lipoprotein containing a signal peptide (SP) for the Sec pathway to be secreted and a lipobox motif to be attached to the membrane via lipid anchoring. Our results indicate that Cpi1 is translocated and localized to the outer membrane cell surface. (1) We hypothesize that PpCpi1 can protect surface-localized proteins from cleavage by inhibiting plant apoplastic PLCPs. Fragments released from the surface-localized proteins could be recognized by plant PRR receptors as MAMPs and trigger PTI responses thus reducing colonization. (2) Cpi1 is localized in OMVs that might be protected from PLCP cleavage. OMVs cargo might help in the establishment of root colonization by *Pseudomonas* and other SynCom members (public good). Dashes lines indicate unknown / not confirmed pathways. Created in BioRender. Doehlemann, G. (2026) https://BioRender.com/rlmkc9o.

Cpi1 is particularly important for bacterial establishment in the meristematic, transition, and elongation zones of *A. thaliana* roots. These regions are characterized by high exudation of primary and secondary metabolites that act as chemoattractants (Canarini *et al*., 2019; Haichar *et al*., 2014; Hale & Moore, 1980; Jones *et al*., 2009), with vasculature-derived glutamine serving as a major bacterial chemoattractant and promoter of proliferation (Tsai *et al*., 2025). Furthermore, the elongation zone might be particularly susceptible to bacterial invasion due to the ongoing remodeling of the thin cell wall and the absence of fully developed endodermal barriers (Lü *et al*., 2022; Somssich *et al*., 2016; Tsai *et al*., 2023). Consistent with this, microbe perception, immune signaling and defense responses are enriched in these root regions (Emonet *et al*., 2021; Millet *et al*., 2010). For example, flg22 or peptidoglycan treatment induces strong expression of MAMP-responsive genes, camalexin secretion and callose deposition in the elongation zone of *A. thaliana* roots (Millet *et al*., 2010). Together, these observations suggest that the root tip and elongation zone are nutrient-rich yet immunologically active niches, where successful colonization likely requires balancing nutrient acquisition with suppression or evasion of host immunity (Bulgarelli *et al*., 2013; Dennis *et al*., 2010; Zhalnina *et al*., 2018). In this context, Cpi1-mediated modulation of host PLCP activity may provide an adaptive advantage by facilitating establishment of *P. putida* within these densely colonized root regions, where host immunity and microbial competition likely intersect (Hacquard *et al*., 2017; Venturi & Keel, 2016).

The observed alterations in the synthetic community in the absence of Cpi1 suggest that its activity is not limited to the producing strain. Consistent with this idea, Cpi1 was detected in outer membrane vesicles (OMVs), which are known to mediate the extracellular delivery of proteins and other cargos. This raises the possibility that Cpi1 may function beyond the cell of origin, potentially acting as a public good that influences neighboring microbes within the community (Fig. 6). OMVs are known to carry a diverse array of biomolecules including proteins, lipids and small molecules that contribute to bacterial virulence and intercellular communication (Orench-Rivera & Kuehn, 2016; Rybak & Robatzek, 2019; Schwechheimer & Kuehn, 2015). Consistent with our findings, OMVs from *P. syringae* pv. *tomato* have been reported to be enriched in virulence factors, including Cpi1, under apoplast-mimicking conditions, suggesting that OMV-mediated transport may facilitate the delivery of protease inhibitors to the plant interface (McMillan & Kuehn, 2023; Tran *et al*., 2022). In addition, Cpi1 orthologs have been identified in OMVs from *P. putida* KT2440 and *P. aeruginosa* PAO1 (Ballok *et al*., 2014; Choi *et al*., 2014; Park *et al*., 2015), supporting the association of chagasin-like inhibitors with OMVs in *Pseudomonas*. OMV-associated cargo has been shown to benefit not only the producing cells but also neighboring bacteria, supporting the idea that OMVs can mediate community-level effects (Valguarnera *et al*., 2018). In this context, OMV-associated Cpi1 may contribute to shaping the root-associated microbial community, consistent with the role of *P. putida* AA7 as a keystone member of the maize root microbiome (Niu *et al*., 2017). However, the mechanisms underlying these effects remain to be determined.

In summary, our work identifies Cpi1 as a surface-exposed inhibitor of plant proteases that promotes early root colonization by the commensal bacterium *P. putida*. These findings highlight the interplay between microbial colonization strategies and host immunity and raise the possibility that Cpi1 additionally contributes to shaping microbial community structure. Collectively, this work expands our understanding of how rhizosphere-associated bacteria successfully colonize plant roots.

## Material and Methods

### Plant material and growth conditions

*Nicotiana benthamiana* plants were grown for six weeks at 23°C for 16h of light and at 20°C for eight hours of darkness with 30 % to 40 % humidity. *Z. mays* cultivar B73 seeds were washed in 70 % Ethanol for 5 min followed by sterilization with 1.2 % sodium hypochlorite for 15 min and three times washes with sterile Milli-Q water. Seeds were germinated in the dark at RT for three days. Sterilized 50 ml syringes without pistil were filled with sterilized soil:seramis granulate mixture (2:1) and one germinated seed was planted. Maize plants (cultivar B73 or Golden Bantham) grew 16h with light at 28°C and for 8 h in darkness at 22°C with 40 % humidity. *A. thaliana* seeds were sterilized for four hours with chlorine gas and placed into RootChip8NF. The chips were positioned with an angle of 45° into a plant growth cabinet with side illumination with 16 h light and 21 °C.

### Maize SynCom experiment and amplicon sequencing

SynCom strains and Pseudomonas putida variants were grown overnight in dYT at 28 °C with shaking. Two synthetic communities were assembled, each containing six of seven maize root SynCom members plus either *P. putida* WT or *P. putida* Δ*cpi1* (final OD₆₀₀ = 0.0001 in 10 mM MgCl₂). One day after seed germination in soil, plants were inoculated with 10 mL of the respective SynCom suspension. Seedlings were harvested at 3 h, 7 dpi, and 14 dpi, washed twice by vortexing in sterile MilliQ water (10 s, ∼10 mL), and roots were excised and flash-frozen in liquid nitrogen.

Root tissue was ground to a fine powder, and ∼500 mg aliquots were stored at −80 °C. DNA was extracted using the FastDNA Spin Kit for Soil (MP Biomedicals) according to the manufacturer’s instructions. The V5–V7 region of the 16S rRNA gene was amplified using primers 799F (AACMGGATTAGATACCCKG) and 1192R (ACGTCATCCCCACCTTCC) following established protocols (Durán *et al*., 2018; Harbort *et al*., 2020). Libraries were prepared using the Novoseq pipeline and sequenced by Novogene UK Ltd. Reads were demultiplexed and primer-trimmed using cutadapt in QIIME2 (maximum error rate 0.25, 10 nt overlap) (Bolyen *et al*., 2019). Sequences were mapped to SynCom reference 16S rRNA gene sequences using BWA-MEM with a minimum mapping quality of 55. Read counts were normalized to 16S rRNA copy number obtained from whole genome sequencing, and relative abundances of each bacterial group were calculated as the proportion of reads per sample. Community dissimilarity was assessed using Bray–Curtis distances followed by principal coordinate analysis (PCoA). Goodness of fit and PCoA was calculated using the R package vegan (Oksanen *et al*., 2018). Two sided-sided t-test was performed comparing SynCom members plus either *P. putida* WT or *P. putida* Δ*cpi1* for each genus and timepoint.

### *A. thaliana* RootChip experiments

Root colonization was analyzed using the RootChip8NF microfluidic device, a modified version of the RootChip8S system adapted for 3D-printed molds and optimized for horizontal and vertical microscopy. Sterile Arabidopsis thaliana Col-0 seedlings were germinated on MS agar and transferred into RootChip8NF devices three days after germination. *Pseudomonas* strains were grown overnight in LB supplemented with gentamycin (34 µg ml⁻¹), diluted to OD₆₀₀ = 0.02 in Hoagland’s medium, and introduced into chips containing 10-day-old seedlings.

Roots were imaged 1 day post inoculation using a custom-built inverted epifluorescence microscope equipped with a 25× water-immersion objective, automated stage, laser illumination system, and sCMOS camera. Consecutive overlapping fields of view covering up to 5 mm from the root tip were acquired as Z-stacks and processed by background correction, maximum-intensity projection, and automated threshold-based segmentation.

Segmented bacterial objects were mapped relative to a manually defined root midline to determine their position along the root axis. Only objects within 200 µm of the root surface and 4 mm from the root tip were included. Bacterial coverage was quantified in consecutive 200 × 100 µm bins along the root and analyzed in R using generalized additive models (GAMs) with distance from the root tip as a smooth term and imaging session and image identity as random effects. Position-dependent autofluorescence background was subtracted before modeling. Statistical differences from WT were determined based on non-overlapping 95% confidence intervals. Root tip colonization assays were normalized to the median WT control within each imaging session and analyzed using two-sided Wilcoxon rank-sum tests with Benjamini–Hochberg correction.

### Cloning of recombinant Cpi1 and PLCP constructs

Vectors for heterologous expression of maize PLCPs in *N. benthamiana* were generated using Golden Gate Modular Cloning (MoClo) with standard plant expression modules (Engler *et al*., 2014; Weber *et al*., 2011). Maize PLCP genes for CP1A and RD19A-like were amplified from *Zea mays* B73 cDNA and cloned into Level 0 or Level 1 vectors. Internal BsaI sites in THI1-like were removed by site-directed mutagenesis. For recombinant protein expression in *E. coli*, PpCpi1 was cloned into the pOPIN-GG system (Bentham *et al*., 2021) to generate N-terminal His-SUMO-tagged constructs. All cloning reactions were performed according to established MoClo and pOPIN-GG protocols. Primer sequences used in this study are listed in Supplementary Table 1.

### Transient expression of PLCPs in *N. benthamiana* leaves

Transient expression of PLCPs and collection of apoplastic fluid was performed as described by (Mueller *et al*., 2013). A 1:1 mix of *A. tumefaciens* cultures carrying plasmids for the expression of PLCPs or P19 plasmid (OD_600_ = 1) in 10 mM MgCl_2_ buffer was infiltrated into 4-6 weeks old *N. benthamiana* leaves.

### Cloning and plasmid construction

sfGFP, mCherry, Flag-tagged Cpi1, and control constructs were generated by PCR amplification and restriction-based cloning using standard molecular biology approaches. Constitutive expression vectors were generated by replacing the inducible pTac promoter of pJT’Tmcs (Verhoef *et al*., 2010) with the constitutive pNm promoter (Labes *et al*., 1990). Cpi1 fusion constructs containing C-terminal sfGFP, mCherry, or Flag tags were generated and sequence verified following transformation into E. coli. Constructs were subsequently introduced into *P. putida* WT, Δ*cpi1*, WT-sfGFP, and Δ*cpi1*-sfGFP strains. Additional PsCpi1 and PsCpi1C18A constructs for expression in *P. syringae* DC3000 were generated similarly using pML123-based vectors. Primer sequences used in this study are listed in Supplementary Table 1.

### Bacterial transformation

*E. coli* was transformed by heat shock, whereas *P. putida, P. syringae, and A. tumefaciens* were transformed by electroporation as described previously (Huang & Wilks, 2017; Schulze Hüynck *et al*., 2019).

### Generation of Pseudomonas mutant and fluorescent strains

Deletion and genomic integration constructs were generated using Gibson Assembly into pK19mobsacB-based allelic exchange vectors (Huang & Wilks, 2017; Schäfer *et al*., 1994). The *cpi1* deletion mutant, sfGFP knock-in strains, and complementation strains were generated by two-step allelic exchange at the attTn7 site downstream of *glmS* (Matsumoto *et al*., 2022). Domain deletion strains were generated similarly. Mutants were selected on sucrose-containing medium and verified by PCR and sequencing.

### Isolation of bacterial outer membrane vesicles (OMVs)

OMVs from *P. putida* WT and Cpi1-overexpressing strains were isolated using either a modified ExoBacteria™ OMV Isolation Kit protocol (SBI, EXOBAC100A-1) or density gradient ultracentrifugation. For kit-based isolation, cultures were grown in 500 mL dYT (28 °C, shaking) to OD₆₀₀ 1.5–2.0. Cells were removed by centrifugation (5,000 × g, 15 min, 4 °C) and supernatants filtered (0.22 μm). OMVs were precipitated with ammonium sulfate to 70% saturation, incubated overnight at 4 °C, and pelleted (10,000 × g, 30 min, 4 °C). Pellets were resuspended in PBS, incubated with resin (1 mL, 30 min, 4 °C), and purified according to the manufacturer’s protocol. For ultracentrifugation-based isolation, cultures were grown in 150 mL dYT to OD₆₀₀ 2.0. After removal of cells (7,000 × g, 30 min, 4 °C) and filtration (0.22 μm), supernatants were ultracentrifuged (150,000 × g, 60 min, 4 °C). Pellets were resuspended in 50% OptiPrep and separated on 15–40% OptiPrep gradients. After centrifugation (100,000 × g, 6 h, 4 °C), OMV-containing fractions (identified by FM4-64 staining) were pooled, diluted in PBS, and pelleted again (150,000 × g, 2 h, 4 °C). Final OMVs were resuspended in PBS.

### Transmission Electron Microscopy (TEM) and OMV quantification

To observe OMVs by transmission electron microscopy, 10 µL of isolated/purified OMVs (Optipräp 28.01.2026 – WT 30% - Cpi1-Flag 25%) were incubated onto carbon-coated copper grids (CF400-Cu-50, EMS, Morgantown, PA, USA) previously glow-discharged in a Quorum Gloqube plus (Quorum Technologies, Laughton, East Sussex, UK). Samples were allowed to settle for 15 minutes before washing with de-mineralized water and negative staining with 2% aqueous uranyl acetate, twice for one minute. After drying overnight in an oven at 40 °C, grids were examined with a Hitachi HT7800 TEM (operating at 100 kV) equipped with an EMSIS Xarosa digital camera. Representative micrographs of OMVs were taken at the same magnification (x40k). Vesicle segmentation was then realized using the Cellpose Cyto3 model (Stringer & Pachitariu, 2025) after removal of edge artifacts. The diameter of OMVs was quantified from ten micrographs per genotype (WT, n = 936; Cpi1-Flag, n = 521).

### Growth curves of bacteria

Growth curve measurements were performed using the Tecan plate reader with a starting OD_600_ of 0.01 in 100 µl dYT medium in 96 well plates. The plates were incubated in the plate reader at 28°C with a constant shaking of 3.5 rpm and OD_600_ was measured for 24 hours.

### Fluorescence imaging of bacteria

*P. putida* WT-sfG and Cpi1-mCherry strains were diluted and washed twice in PBS and 10 µl of each sample was fixed by short heat application to a microscope slide. The samples were observed with an 100x oil immersion objective (NA: 1.45) using a Nikon Eclipse Ti operating system and a Hamamatsu orca-flash4.OLT digital camera. For sfGFP and mCherry excitation 488 nm and 543 nm were used, respectively. The emission of sfGFP and mCherry were measured between 500 nm to 535 nm and 575 and 650 nm for mCherry, respectively. The focus was based on the z-position with strongest GFP signal. Fluorescence intensities of sfGFP and mCherry throughout the cells were measured: A line (white line in graphs) was drawn with ImageJ orthogonal to the longer side of the cell in the sfGFP and mCherry images and a plot profile of the intensity was measured. Within each profile the intensities were normalized to the minimum (0%) and maximum intensity (100%) of each profile. Next, the center position of the cell was determined as the position with the maximum sfGFP signal for each pair of sfGFP-mCherry profiles. A paired two-sided t-test was performed for the signals of mCherry and sfGFP at approx. -0.52 μm, 0 μm, and 0.52 μm distance from the center using the t-test function in R.

### Heterologous protein expression and purification

His-SUMO-tagged *P. putida* Cpi1 was heterologously expressed as described for the pOPIN system (Bentham *et al*., 2021). After heterologous protein production, cells were resuspended in Ni-NTA lysis buffer (50 mM NaPO_4_ pH 8, 300 mM NaCl,10 mM Imidazol, 0.01% (v/v) Benzoase, 1X cOmplete protease inhibitor cocktail) and sonicated (three cycles of 5 min with 0.5s pulse and 40% amplitude). The soluble fraction was obtained using 25 min centrifugation at 20000 rpm at 4°C. His-tagged proteins were purified using a 1 ml HisTrap FF Crude column on a ÄKTA start system equilibrated with Ni-NTA binding buffer (20 mM NaPO4 pH 7.4, 500 mM NaCl, 25 mM Imidazol) using the affinity chromatography standard program and were eluted with the Ni-NTA elution buffer (20 mM NaPO4 pH 7.4, 500 mM NaCl, 500 mM Imidazol). To remove the SUMO solubility tag, the purified His-SUMO-PpCpi1 was incubated with 250 µl precision protease in cleavage buffer (50 mM Tris HCl, 100 mM NaCl, 1 mM EDTA, 1 mM DTT, pH 7.5) overnight at 4°C. PpCpi1 was purified via size-exclusion chromatography on a HiLoad 16/600 Superdex 75 pg column using an ÄKTA pure system using 1.5 CV of the equilibration buffer (50 mM Tris-HCl 150 mM NaCl pH7). The protein concentration was determined using a Bradford Assay with the 2X ROTI Quant reagent according to the manufacturer’s manual.

### Gel electrophoresis and western blot analysis

Proteins were denatured in 1× SDS loading buffer at 98 °C for 5 min and separated on 15% SDS–PAGE gels. Proteins were transferred onto 0.2 µm PVDF membranes using the Trans-Blot Turbo Transfer System (Bio-Rad). Membranes were blocked in 3% milk in TBS-T and incubated with primary antibodies against Flag (Sigma, A8592), GFP (Roche, 47859600), IcdH (Merck, ABS2090), OprF (Invitrogen, PA5-117553), or streptavidin-HRP (Sigma, S2438). HRP-conjugated anti-mouse or anti-rabbit secondary antibodies (Cell Signaling) were used where appropriate. Signals were detected using SuperSignal West Pico PLUS or Femto substrates (Thermo Fisher) and visualized with a ChemiDoc imaging system. Western blot analysis of PsCpi1 was performed as described previously (Shindo *et al*., 2016).

### Activity-based protein profiling (ABPP) and substrate cleavage assay

Activity-based protein profiling (ABPP) was performed as described by (Schulze Hüynck *et al*., 2019) with 0.6 µM MV201 (Richau *et al*., 2012), DCG04-Cy5 (Stolze *et al*., 2012), or DCG04 (Greenbaum *et al*., 2000). MV201 and DCG-04-Cy5 were kindly provided by Hermen Overkleeft (Leiden University). The substrate cleavage assay was performed as described by (Schulze Hüynck *et al*., 2019) using the synthetic substrate Z-LR-AMC. IC_50_ and statistics were calculated using the Hill-function in GraphPad Prism (Neubig *et al*., 2003).

### Cell fractionation

Cell fractionation was performed largely as described previously (Molloy *et al*., 1998; Paredes-Osses & Hardie, 2013). Briefly, fresh Pseudomonas putida cultures (OD_600_ = 1) were disrupted using a Constant Systems cell disruptor (2.5 kBar). Cytosolic and membrane fractions were separated by ultracentrifugation (183,000 g, 60 min, 4 °C). Periplasmic fractions were isolated by osmotic shock, followed by freeze–thaw lysis to obtain a second cytosolic fraction. Inclusion bodies were solubilized in 8 M urea, 40 mM Tris pH 9.5, 100 mM DTT, and membrane pellets were resuspended in 1% SDS.

### Biotin labeling

Surface proteins of *P. putida* Cpi1-Flag cells (OD_600_ = 0.3) were labeled using EZ-Link Sulfo-NHS-Biotin (Thermo Fisher) according to the manufacturer’s instructions. Cells were lysed in membrane protein buffer (50 mM Tris-HCl pH 8, 150 mM NaCl, 1% Triton X-100, 0.5% sodium deoxycholate, protease inhibitors) by sonication on ice. Biotinylated proteins were captured using streptavidin beads, washed, and eluted in SDS loading buffer at 95 °C for 10 min.

### Bioinformatic analysis

SynCom genomes were annotated using dFast (Tanizawa *et al*., 2016, 2018). BLASTp searches were performed with BLAST+ (Camacho *et al*., 2009) using MEROPS family C1 PLCP sequences as queries (E-value cutoff: 0.001). DIAMOND BLASTp searches against complete proteomes from the Pseudomonas Genome Database were performed using PpCpi1 as query (E-value cutoff: 1 × 10−4, query coverage ≥40%, identity ≥40%) (Buchfink *et al*., 2015; Winsor *et al*., 2016). Signal peptides were predicted using SignalP6 (Teufel *et al*., 2022). Multiple sequence alignments were generated with CLUSTAL Omega (Sievers *et al*., 2011) and visualized with MView

(Brown *et al*., 1998). Chagasin motifs were identified by regular expression searches and visualized using ggVennDiagram and ggseqlogo (Gao *et al*., 2021, 2024; Wagih, 2017). Protein structures were predicted with AlphaFold3 (Abramson *et al*., 2024) and analyzed using PyMOL.

### Quantitative real-time PCR

Quantitative real-time PCR (qPCR) was performed using Promega GoTaq qPCR Master Mix on a CFX Connect Real-Time System (Bio-Rad; SYBR mode). Reactions were run in technical duplicates. Cycling conditions were 95 °C for 2 min, followed by 45 cycles of 95 °C for 30 s, 61 °C for 30 s, and 72 °C for 30 s, with a melt curve from 65–95 °C. Ct values were calculated using Bio-Rad CFX Manager software. Relative *cpi1* expression was quantified using the 2−ΔΔCt method with *ropD* as the reference gene (Livak & Schmittgen, 2001). Ct values >35 were considered background; for visualization on log-scale plots, these were set to 0.0001. Primer sequences used in this study are listed in Supplementary Table 1.

## Data availability

Source data are provided in this paper. All data supporting the findings of this study that are not directly available within the paper (and its supplementary data) will be upon reasonable request available from the corresponding authors (GD, JM). All scripts are deposited on GitHub (https://github.com/dmoser1/PpCpi1_preprint.git).

## Accession codes

Chagasin: Q966X9, PpCpi1: BUQ73_RS03925, PaCpi1: PA0778, PfCpi1: Pfl01_4580, PsCip1/PsCpi1: PSPTO_4211, RD19A-like: Zm00001eb240700, CP1A: Zm00001eb068400.

## Acknowledgments

We are grateful to Jan Muelhoefer, Dorrie Wamser, and Sabrina Egli for their technical support. We would like to thank Dr. Noah Kürtös and Prof. Dr. Ryohei Thomas Nakano for their support during the preparation of samples for amplicon sequencing and in the data analysis. We are thankful to Lucie Hansen for establishing the biotin labeling of surface proteins. The pJT’Tmcs vector and the pK19-moB-sacB vector were kindly provided by Dr. Anita Loeschke. We thank Prof. Dr. Hermen Overkleeft for providing MV201 and DCG04-Cy5 for the ABPP experiments. We also want to thank Tomohisa Shimasaki for sharing the protocol for OMV isolation and valuable assistance. We acknowledge support from the Cluster of Excellence on Plant Sciences (CEPLAS) funded by the Deutsche Forschungsgemeinschaft (DFG, German Research Foundation) under Germany’s Excellence Strategy – EXC 2048/1 – project ID 390686111 and a DFG Heisenberg Professorship (GR4559/4-1, to G.G.).

## Author contributions

DM, GD and JMV designed the study. DM performed the bioinformatics analysis. DM, NS and UM performed the cloning, biochemical experiments and fluorescence microcopy. FK, BM and RVDH performed the biochemical characterization of the chagasin motif in *P. syringae*. UN performed electron microscopy and AV quantification of OMVs. DM and CFK performed the *A. thaliana* root colonization experiments and the data analysis together with GG and JMV. DM and LR performed the amplicon sequencing experiment together with TGA and JMV. DM and JMV wrote the manuscript with contributions from all authors.

---

**Supplementary Figure S1:**
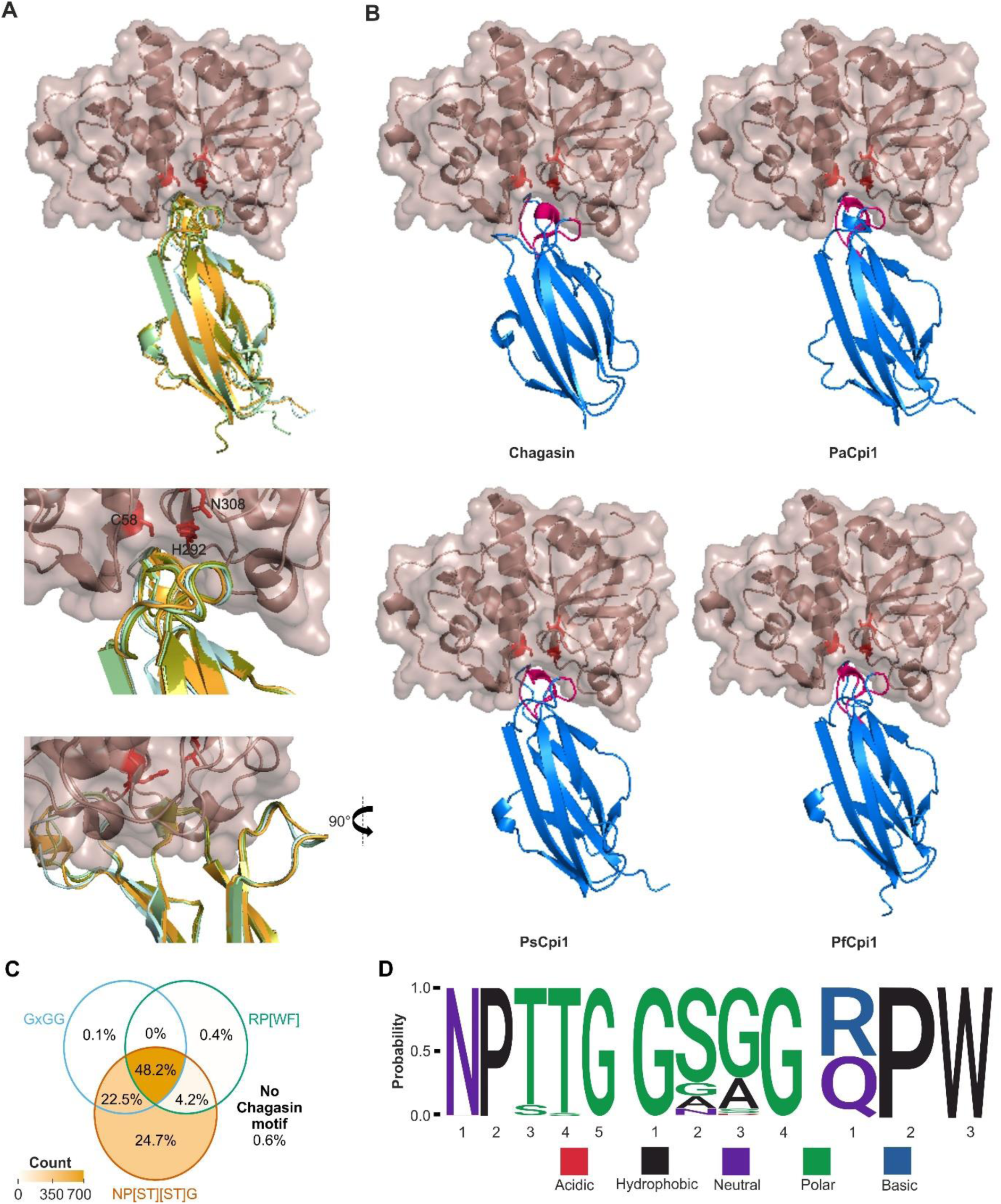
Conserved chagasin-like inhibitors in pseudomonads are predicted to block the PLCP’s active site via their chagasin motif. **(A, top panel)** Chagasin-like protease inhibitors (PpCpi1: light green, PsCpi1: light blue, PfCp1: yellow, PaCpi1: orange) are predicted to inhibit PLCPs (papain, dark red) by blocking the protease active site (red) with their chagasin motifs (pink). *Pseudomonas* chagasin-like inhibitors were modelled with AlphaFold3 (Abramson *et al*., 2024) without signal peptide. The models were aligned to the structure of TcChagasin in a complex with papain (PDB: 3E1Z) (Redzynia *et al*., 2009) and overlaid in PYMOL. **(A, bottom panel)** Zoom on loops with chagasin motifs (pink) blocking active site of papain (red). Catalytic triad is labeled. **(B)** Complex models of the individual chagasin-like inhibitors (blue) from A in complex with papain (dark red) blocking the protease active site (red) with their chagasin motifs (pink). **(C)** 1527 complete *Pseudomonas* proteomes deposited in the *Pseudomonas* Genome Database (Winsor *et al*., 2016) were screened with a DIAMOND BLASTp search (Buchfink *et al*., 2015) for orthologs of PpCpi1 (E < 0.0001). PpCpi1-orthologs were aligned via CLUSTAL (Sievers *et al*., 2011) and tested for presence or absence of the original chagasin motifs (NP[ST][ST]G, GxGG, and RP[WF]) (dos Reis *et al*., 2008; Shindo *et al*., 2016). Presence and absence were visualized using a Venn diagram (Gao *et al*., 2021, 2024) to illustrate the overlap of different chagasin motifs. **(D)** Motif logos of the chagasin motifs identified in pseudomonads were created using ggseqlogo (Wagih, 2017). Amino acids are colored based on their chemical properties.

**Supplementary Figure S2:**
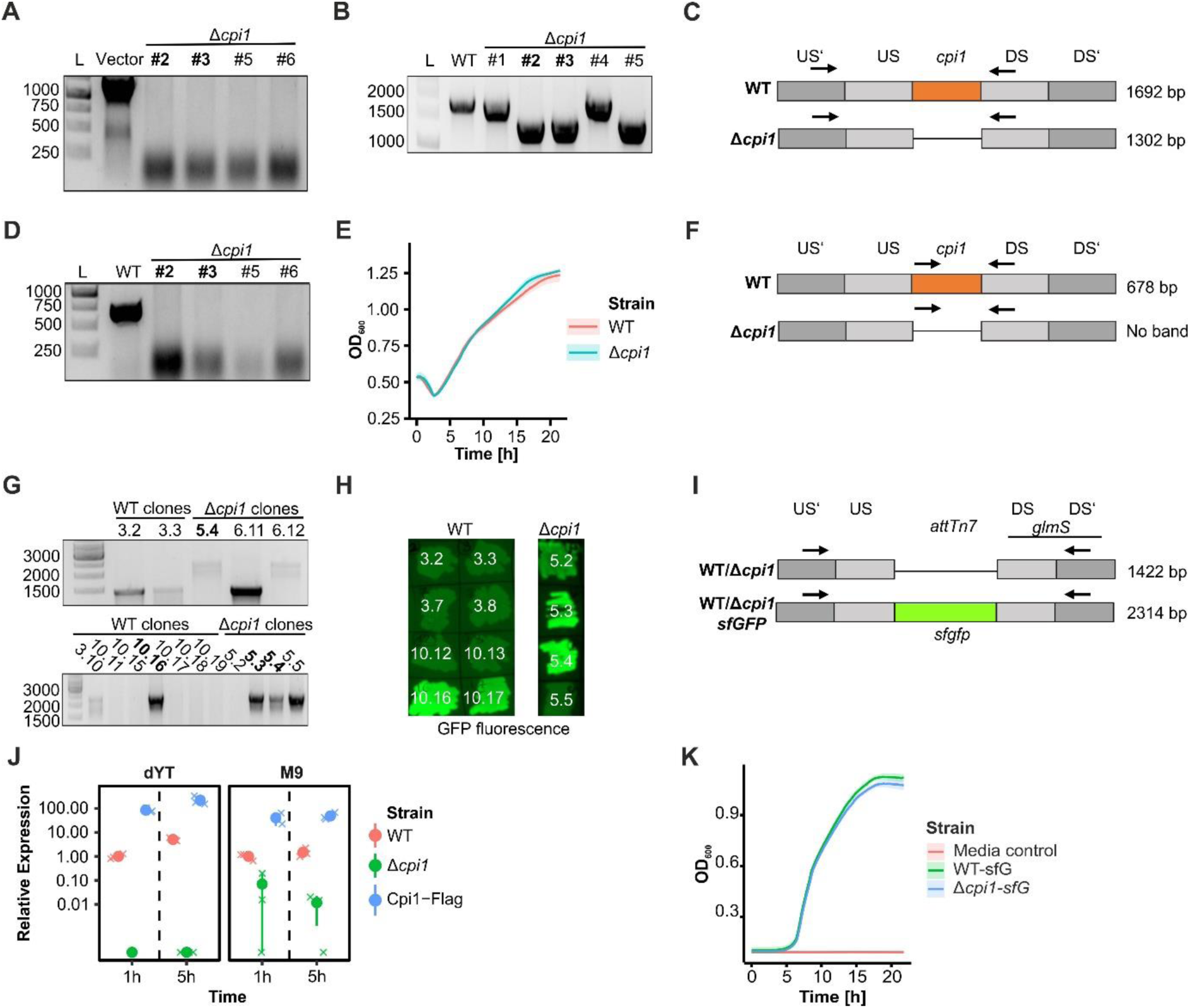
Generation and validation of genomic knockout Δ*cpi1* and integration of *sfgfp* into *P. putida* strains. The genomic deletion of *cpi1* was tested using multiple rounds of colony PCR. Two clonesΔ*cpi1*#2 and Δ*cpi1*#3 were confirmed via sequencing. **(A)** The deletion of *cpi1* was furthermore tested via PCR with primers binding in the *cpi1* and DS sequence. The absence of a band at 678 bp suggests deletion of *cpi1*. **(B, C)** Deletion of *cpi1* between upstream sequence (US) and downstream (DS) sequence was tested via PCR with primers binding in the US’ and DS region. A 1302 bp band indicated absence of the *cpi1* sequence. **(D, F)** The loss of the vector was tested with vector specific primers. **(E)** Growth curve of *P. putida* WT and Δ*cpi1*#2 grown in full medium. Tecan plate reader was used to measure the optical density (OD_600_) of strains grown in dYT medium at 28°C with constant shaking for 24 hours. Dark line represents the mean of three (Δ*cpi1*) / five (WT) replicates. Shadows display standard deviation of the measurement. **(G, I)** The genomic integration of *sfgfp* was tested using colony PCR and sequencing confirmed genomic integration of *sfgfp* in WT_sfG clone #10.16, Δ*cpi1_*sfG clones #5.3 and #5.4. A 2314 bp band indicated the presence of the *sfgfp* sequence in the attTn7 locus upstream of the *glmS* gene. **(H)** GFP fluorescence of clones was evaluated using ChemiDoc Imager. **(J)** RNA was isolated from WT, Δ*cpi1 and* Δ*cpi1*_*cpi1-flag* grown/incubated for one or five hours in dYT or M9 medium. Gene expression was determined by qPCR on cDNA using *cpi1*- as well as reference gene *ropD*-specific primers (Ct > 35 were removed). Relative expression was calculated using 2^-ΔΔCT^ method (Livak & Schmittgen, 2001) and normalized to the value of WT set to 1. Point range displays the average of three replicates (point) with the standard error (line). Crosses represent individual replicates. Not detected or values derived from Ct>35 values were set to 0.0001 for plotting with log-scale. This experiment was performed with three independent replicates. **(K)** Growth curve of WT-sfG and Δ*cpi1-sfG* grown in full medium. Tecan plate reader was used to measure the optical density (OD_600_) of strains grown in dYT medium at 28°C with constant shaking for 24 hours. Dark line represents the mean of six replicates. Shadows display standard deviation of the measurement.

**Supplementary Figure S3:**
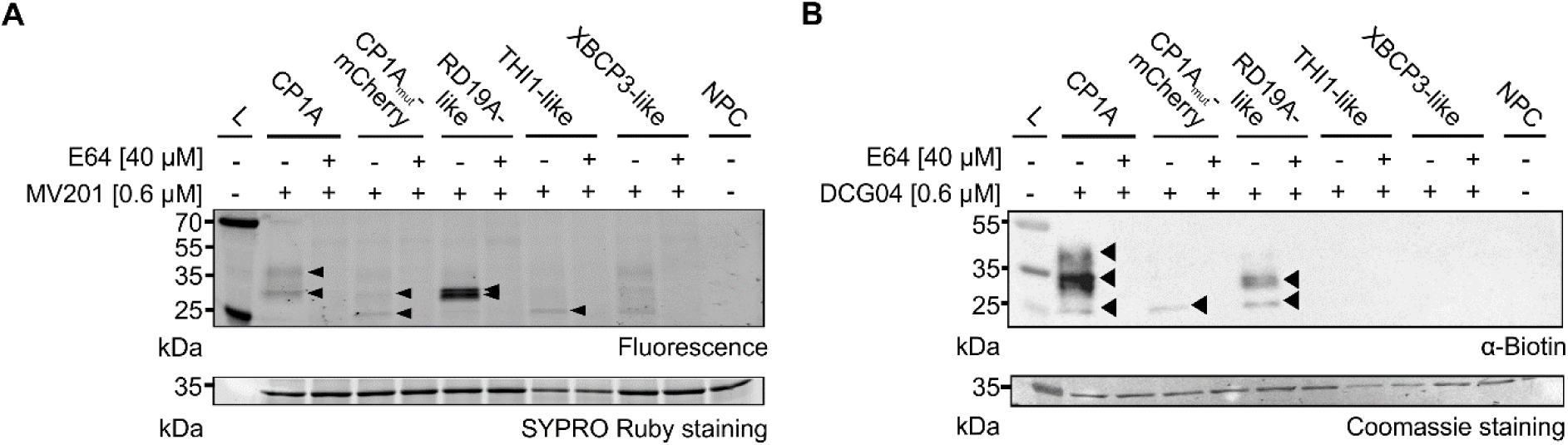
Heterologous expression of maize root PLCPs in apoplastic fluid of *Nicotiana benthamiana* via agrobacterium-mediated transformation. Maize SA-specific root PLCPs (RD19A-like, THI1-like, XBCP3-like) (Schulze Hüynck *et al*., 2019) as well as immune-related CP1A (Schulze Hüynck, 2019; van der Linde *et al*., 2012) and inactive CP1A_mut_-mCherry (Schulze Hüynck, 2019) were heterologously expressed in *N. benthamiana* leaves via agrobacterium-mediated transformation and were secreted into the leaf apoplast using the NbPR1secretion signal. Leaf apoplastic fluid (LAF) was isolated and used for activity-based protein profiling using the PLCP-specific fluorescent probe MV201 (Richau *et al*., 2012) in **(A)** or the biotinylated DCG-04 probe (Greenbaum *et al*., 2000) in **(B)**. PLCP inhibitor E64 (Hanada *et al*., 1978) was used as a control for PLCP activity. LAFs were mixed and incubated without E64 (-) nor MV201(-) and used as a none-probe control (NPC). SYPRO Ruby stating or Coomassie was used as a loading control.

**Supplementary Figure S4:**
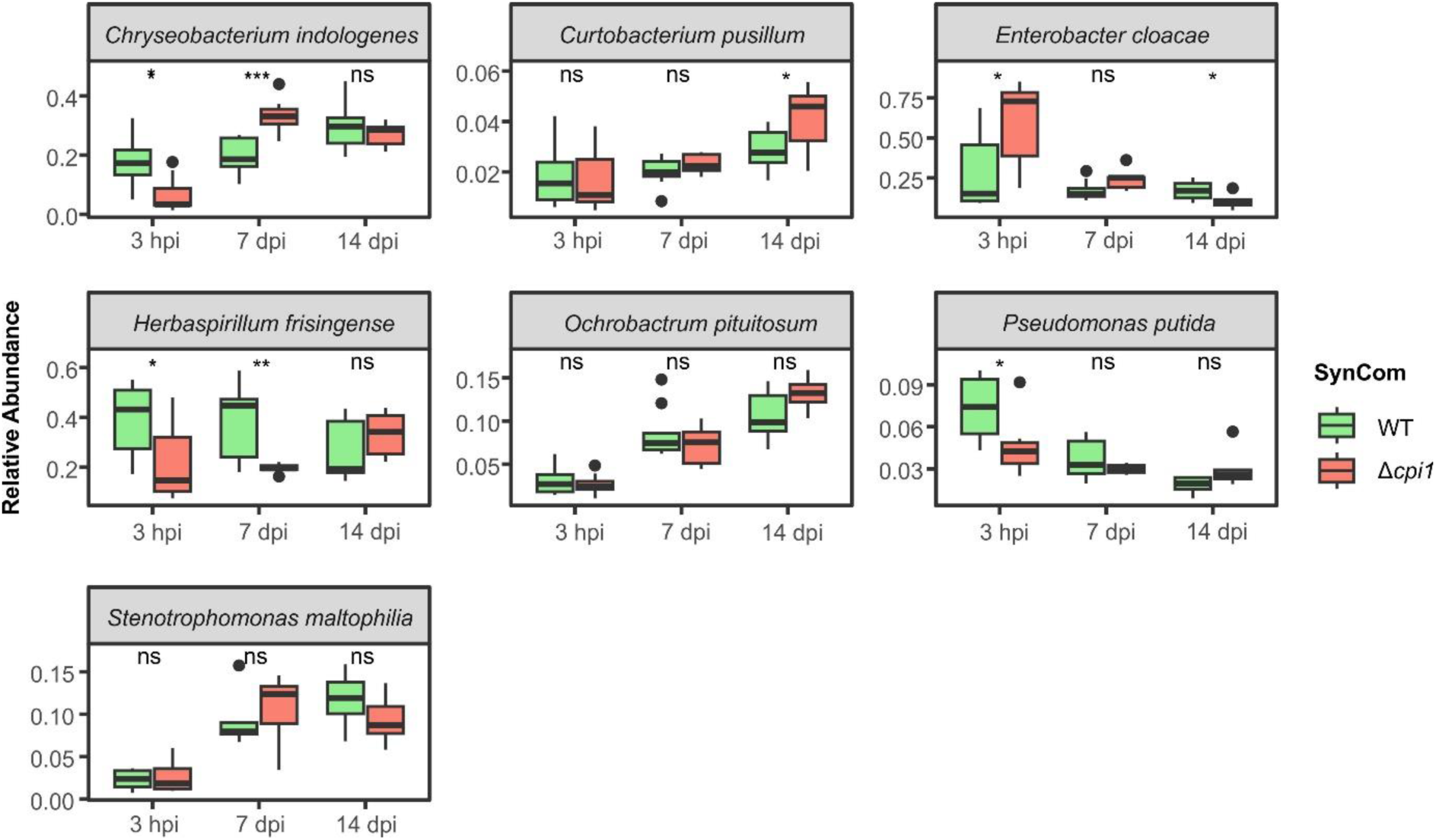
Cpi1 modulates the structure of the synthetic community members (SynCom). Maize plants were grown in sterile soil containing the six SynCom members with either *P. putida* WT or Δ*cpi1* (inoculated to OD_600_ of 0.0001). Maize roots were collected at 3 hpi, 7 dpi and 14 dpi from maize roots, washed several times by vortexing in water, and samples were frozen in liquid nitrogen. DNA was isolated and 16S sequence was amplified via PCR, barcoded and sent for amplicon sequencing. Reads were aligned to the SynCom members 16S RNA sequences and quantified. This experiment was performed once with eight biological replicates per timepoint and SynCom. The abundance of all SynCom members was quantified based on the aligned reads and normalized to the total amount of reads in a sample. A two-sided t-test was performed (P<0.05: *; P<0.01: **; P<0.001: ***; P>0.05: ns).

**Supplementary Figure S5:**
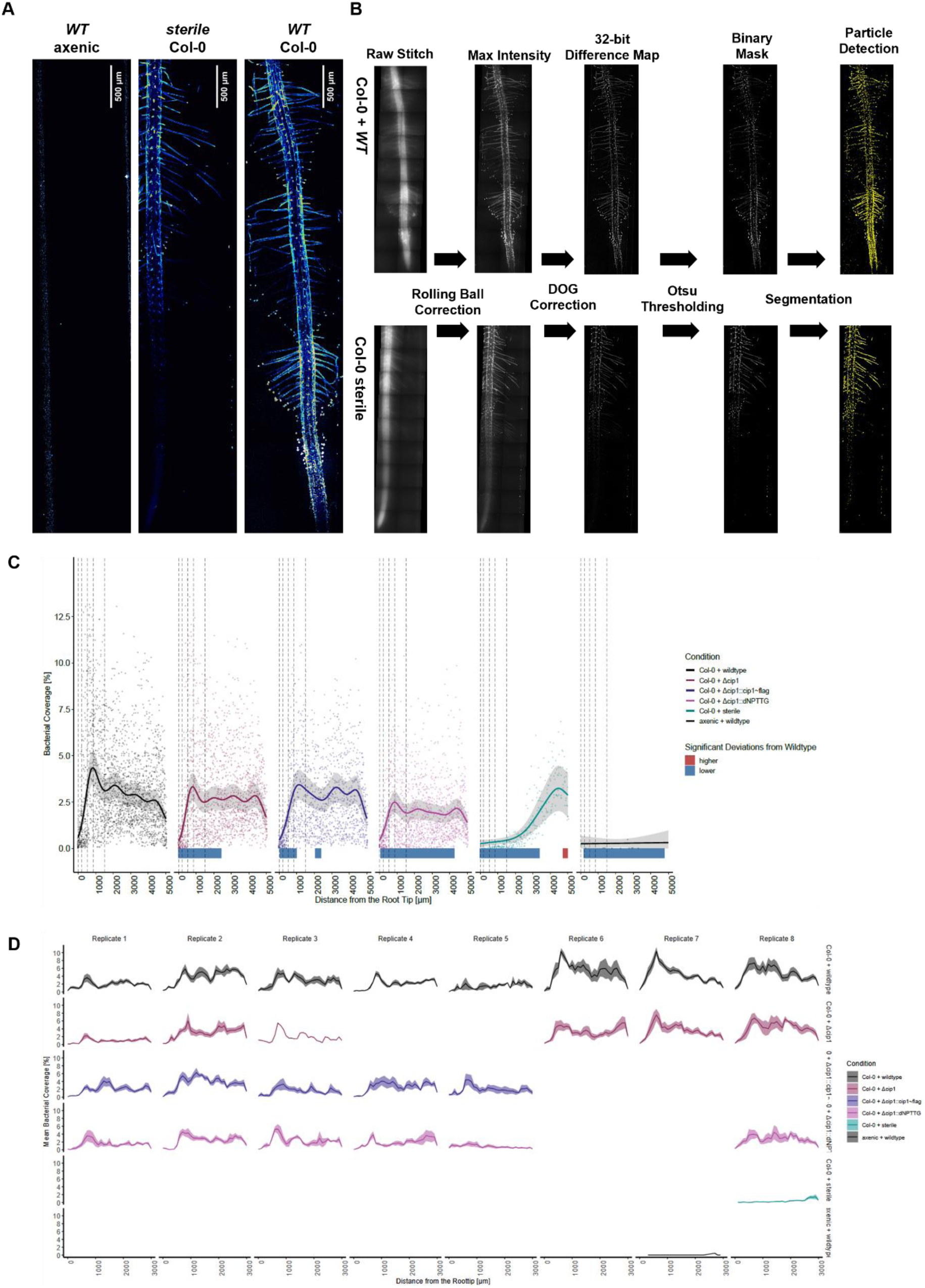
Imaging controls and within-replicate variation of root colonization on *A. thaliana* Col-0. (A) Fluorescence microscopy of colonized A. thaliana Col-0 roots. Maximum-intensity projections of *P. putida* WT-sfG colonizing Arabidopsis thaliana Col-0 roots. Shown are: (i) axenic chip control without a root, (ii) un-inoculated Col-0 roots, and (iii) wild-type–inoculated Col-0 roots. All images are displayed using identical scaling and the “royal” LUT. **(B) Segmentation and background correction workflow.** The upper panel shows an inoculated root; the lower panel shows a sterile, non-inoculated control. Raw images represent summed-intensity projections of the full stitched Z-stack (18 FOVs, ∼5600 µm root length). Background subtraction was performed using a rolling-ball algorithm (radius = 7.5 µm), followed by subtraction of a Gaussian-blurred image (radius = 15 µm) to generate a difference-of-Gaussian (DoG) local background correction. Otsu auto-thresholding produced an 8-bit binary mask (0/255). Particles meeting a circularity threshold >0.3 were retained, excluding linear cell-wall autofluorescence; outlines of detected particles are shown in yellow. **(C) Generalized Additive Modelling of the raw coverage by fluorescent clusters.** Fluorescence intensity was used to segment fluorescent clusters (single cells and aggregates; minimal size ∼3.5 µm²). The segmented bacterial area was summed within 200 × 100 µm windows centered on the root midline and normalized to the bin area to obtain bacterial surface coverage. Colonization profiles along the root were modeled using generalized additive models (GAMs) with bacterial coverage as the response and distance from the root tip as a smooth term. Condition-specific smooths were fitted while accounting for imaging date and image identity as random effects. The solid line shows the fitted GAM and the shaded region indicates the standard error. Individual measurements are shown as jittered points. Regions where colonization significantly differs from the wild type are indicated below the plot (blue = lower, red = higher), based on non-overlapping model confidence intervals. **(D) Bacterial coverage per replicate.** For each imaging session (biological replicate), bacterial coverage was averaged across all roots (solid line), with shaded regions indicating the standard error as a function of distance from the root tip. Coverage was computed in 200 × 100 µm (width × height) bins centered on the root midline. Within each bin, the total segmented bacterial area was summed and normalized to the bin area.

