## Supplemental Table 1 for "A membrane-anchored inhibitor of papain-like cysteine proteases promotes *Pseudomonas* root colonization"

Supplementary Table 1. Primers used in this study

| Primer name | Primer sequence (5′→3′) | Use / Target |
| --- | --- | --- |
| 799F | AACMGGATTAGATACCCKG | Amplicon sequencing (V5–V7 region) |
| 1192R | ACGTCATCCCCACCTTCC | Amplicon sequencing (V5–V7 region) |
| CP1A_F | GGTCTCAAATGGCTGCCTCCACCAC | Amplification of maize CP1A PLCP for MoClo cloning |
| CP1A_R | GGTCTCAAAGCTCATGCGCTGCTCTTCATGC | Amplification of maize CP1A PLCP for MoClo cloning |
| RD19A-like_F | TTGAAGACAAAGGTGTCGACGCGGAGGACCCGCTGA | Amplification of maize RD19A-like PLCP for MoClo cloning |
| RD19A-like_R | TTGAAGACAAAAGCCTACTCCTTCGAGGCGTGGACTGCGGACAC | Amplification of maize RD19A-like PLCP for MoClo cloning |
| Pp*cpi1*_MoClo_F | TTGGTCTCAAATGTGCGCCCAGCAGCCG | Amplification of *P. putida* *cpi1* for MoClo cloning |
| Pp*cpi1*_MoClo_R | TTGGTCTCAAAGCTCAGTCGACGCGGATCGC | Amplification of *P. putida* *cpi1* for MoClo cloning |
| sfGFP_KpnI_F | GATCTAGGTACCATGGGTAAAGGAGAAGAAC | Amplification of sfGFP for tagging constructs |
| sfGFP_XbaI_R | GGAGCTCTCGAGTCTAGAATC | Amplification of sfGFP for tagging constructs |
| sfGFP_XhoI_F | GATCTACTCGAGATGGGTAAAGGAGAAGAAC | Amplification of sfGFP for fusion constructs |
| Pp*cpi1*_KpnI_F | GATCTAGGTACCATGACTGCCCCTCGTCTGC | Amplification of *cpi1* for sfGFP fusion constructs |
| Pp*cpi1*_XhoI_R | GATCTACTCGAGGTCGACGCGGATCGCGCAG | Amplification of *cpi1* for sfGFP fusion constructs |
| pNm_RBS_F | ATGATGGAGCTCGGGCGGTTTTATGGACAG | Amplification of constitutive promoter/RBS cassette |
| pNm_RBS_R | ATGATGGGTACCTCCTCCTAAGCTTGGATCC | Amplification of constitutive promoter/RBS cassette |
| Flag_F | CACTCTGTGGTCTCTCGAGGATTATAAGG | Amplification of Flag-tag sequence |
| Flag_R | TGGTCTCTAGATCACTTATCGTC | Amplification of Flag-tag sequence |
| mCherry_F | ATAGGTACCATGGTGAGCAAGGGCGAGGAG | Amplification of mCherry tag |
| mCherry_R | ATACTCGAGGGCAGCGGCAGCATGGTGAGCAAGGGCGAGGAG | Amplification of mCherry tag |
| Cytosolic_mCherry_R | TGTTCTAGATCACTTGTACAGCTCGTCCATGCC | Amplification of cytosolic mCherry |
| Ps*cpi1*_F | ATCGAAGCTTAGGAGGACAGCTATGCAAACGCCCAAGAACATCGT | Amplification of *cpi1* from *P. syringae* DC3000 |
| Ps*cpi1*_R | CGATGAATTCGTTCACCGTGATTGCGCA | Amplification of Ps*cpi1* from *P. syringae* DC3000 |
| GFP_Ps*cpi1*_F | ATCGCTGCAGGTCGACCCGCGGATGAGTAAAGGAGAAGA | Amplification of GFP cassette |
| GFP_Ps*cpi1*_R | CATGGCATGGATGAACTATACAAATAGATCGATTCTAGACTCGAGGAGCTCATGC | Amplification of GFP cassette |
| Ps*cpi1*_C18A_mut_F | GCGCTGCTGACGGCGGCGGCGCAAACGCCCAAGAAC | Site-directed mutagenesis of Ps*cpi1* signal peptide |
| *cpi1*_KO_up_F | ACAGCTATGACATGATTACGCCAAAGGATGGCCCAAGTGC | Amplification of upstream flank for *cpi1* deletion |
| *cpi1*_KO_up_R | TGACCAGCGCTGTGCGTTCAGGTGGCTCCAGAGGCTGACG | Amplification of upstream flank for *cpi1* deletion |
| *cpi1*_KO_down_F | CGTCAGCCTCTGGAGCCACCTGAACGCACAGCGCTGGTCA | Amplification of downstream flank for *cpi1* deletion |
| *cpi1*_KO_down_R | CCGGGTACCGAGCTCGAATTGTTCATCGCCACCACCGAGG | Amplification of downstream flank for *cpi1* deletion |
| attTn7_up_F | CTATGACCATGATTACGCCAACAGCACCGTGGAGAAAACC | Amplification of upstream attTn7 integration flank |
| attTn7_up_R | GCTGTCCATAAAACCGCCCGCTTGTGTTTTACCGGATGG | Amplification of upstream attTn7 integration flank |
| sfGFP_attTn7_F | CGGGCGGTTTTATGGACAGC | Amplification of sfGFP cassette for genomic integration |
| sfGFP_attTn7_R | ATTATTTGTAGAGCTCATCCATG | Amplification of sfGFP cassette for genomic integration |
| attTn7_down_F | GGATGAGCTCTACAAATAATCCCTGGCCCAAAGCCGGGGC | Amplification of downstream attTn7 integration flank |
| attTn7_down_R | CTTGCGGCAGCGTGAAGCTAGGATCTCGGCGGCAGACAGCC | Amplification of downstream attTn7 integration flank |
| sfGFP_check_F | CCTGCAAGTTACGACGCAGC | Colony PCR verification of sfGFP integration |
| sfGFP_check_R | GTTGACCCTGGCTCTGGGC | Colony PCR verification of sfGFP integration |
| *cpi1*_KO_check_F1 | CGGATGCCAAGATCAAGATCG | Colony PCR verification of *cpi1* deletion |
| *cpi1*_KO_check_R1 | GAGGCGAAGTTAAGGCATTGG | Colony PCR verification of *cpi1* deletion |
| *cpi1*_KO_check_F2 | GAACCCAACCGTGGTGATTTC | Colony PCR verification of *cpi1* deletion |
| *cpi1*_KO_check_R2 | GAAGGGCAGGGTAAGGATGTC | Colony PCR verification of *cpi1* deletion |
| *cpi1*_KO_check_R3 | GGCATCCAACGTGCTACAAAG | Colony PCR verification of *cpi1* deletion |
| ropD_qPCR_F | AAGCGCAACAGCAGTCTCGTATC | qPCR reference gene amplification (ropD) |
| ropD_qPCR_R | CATCCGGAGCACTCTCGAATACG | qPCR reference gene amplification (ropD) |
| *cpi1*_qPCR_F | CCCAGCAGCCGAAACAAACC | qPCR amplification of *cpi1* |
| *cpi1*_qPCR_R | TCTGTACCGGGCGTACTTCC | qPCR amplification of *cpi1* |
